# Genetic and Epigenetic Regulation of Skeletal Muscle Ribosome Biogenesis with Exercise

**DOI:** 10.1101/2020.12.14.422642

**Authors:** Vandré C. Figueiredo, Yuan Wen, Björn Alkner, Rodrigo Fernandez-Gonzalo, Jessica Norrbom, Ivan J. Vechetti, Taylor Valentino, C. Brooks Mobley, Gabriel E. Zentner, Charlotte A. Peterson, John J. McCarthy, Kevin A. Murach, Ferdinand von Walden

## Abstract

Ribosomes are the macromolecular engines of protein synthesis. Skeletal muscle ribosome biogenesis is stimulated by exercise, but the contribution of ribosomal DNA (rDNA) copy number and methylation to exercise-induced rDNA transcription is unclear. To investigate the genetic and epigenetic regulation of ribosome biogenesis with exercise, a time course of skeletal muscle biopsies was obtained from 30 participants (18 men and 12 women; 31 ±8 yrs, 25 ±4 kg/m^2^) at rest and 30 min, 3h, 8h, and 24h after acute endurance (n=10, 45 min cycling, 70% VO_2_max) or resistance exercise (n=10, 4 x 7 x 2 exercises); 10 control participants underwent biopsies without exercise. rDNA transcription and dosage were assessed using qPCR and whole genome sequencing. rDNA promoter methylation was investigated using massARRAY EpiTYPER, and global rDNA CpG methylation was assessed using reduced-representation bisulfite sequencing. Ribosome biogenesis and *MYC* transcription were associated with resistance but not endurance exercise, indicating preferential up-regulation during hypertrophic processes. With resistance exercise, ribosome biogenesis was associated with rDNA gene dosage as well as epigenetic changes in enhancer and non-canonical MYC-associated areas in rDNA, but not the promoter. A mouse model of *in vivo* metabolic RNA labeling and genetic myonuclear fluorescent labeling validated the effects of an acute hypertrophic stimulus on ribosome biogenesis and *Myc* transcription, and corroborated rDNA enhancer and Myc-associated methylation alterations specifically in myonuclei. This study provides the first information on skeletal muscle genetic and rDNA gene-wide epigenetic regulation of ribosome biogenesis in response to exercise, revealing novel roles for rDNA dosage and CpG methylation.

**GRAPHICAL ABSTRACT:** 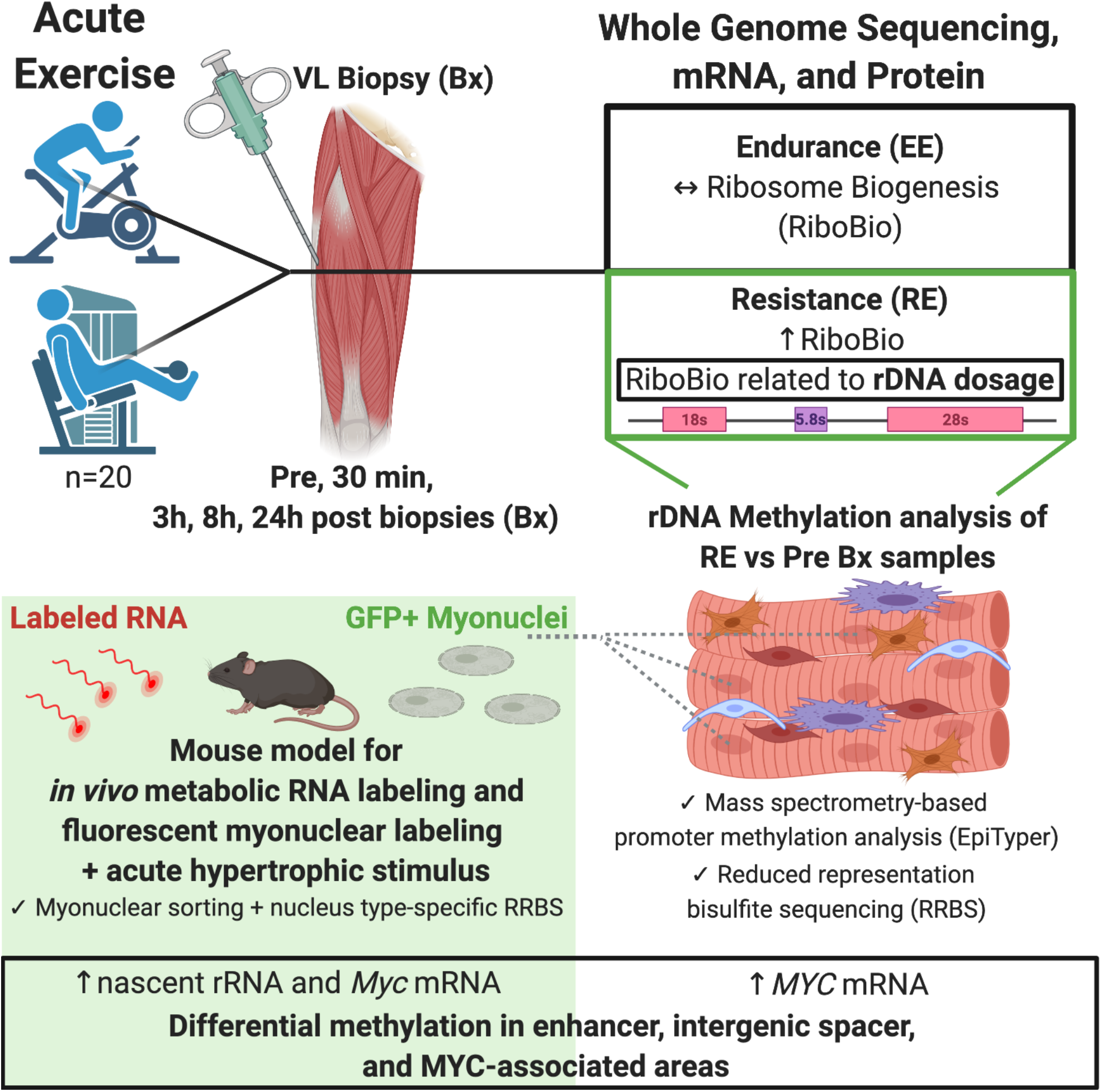

## INTRODUCTION

Ribosomes are the molecular factories responsible for protein synthesis, which is a key process in the long-term adaptive response to exercise in skeletal muscle (Figueiredo, 2019a; McCarthy & Murach, 2019). Ribosome biogenesis is stimulated by muscle loading (Figueiredo & McCarthy, 2019b; Kim *et al*., 2019; von Walden, 2019b), and the magnitude of *de novo* synthesis of ribosomes is proportional to the amount of load-induced adult muscle hypertrophy in both rodents (Nakada *et al*., 2016) and humans (Figueiredo *et al*., 2015; Stec *et al*., 2016; Hammarstrom *et al*., 2020). Mechanical loading rapidly induces RNA Polymerase I (Pol I) transcription of ribosomal DNA (rDNA), as assessed by 45S pre-rRNA transcription levels and accumulation of rRNA (von Walden *et al*., 2012b; Kirby *et al*., 2016; Figueiredo *et al*., 2019b). The production of the long 45S pre-rRNA transcript, which is processed into the mature 18S, 5.8S and 28S rRNAs, is believed to be the rate limiting step of ribosome biogenesis (Moss 2004). Contrary to protein-coding genes that commonly occur in the human genome as two copies, rDNA genes number in the hundreds and vary widely across individuals (Gibbons *et al*., 2015; Malinovskaya *et al*., 2018; Parks *et al*., 2018). While each rDNA locus can participate in the formation of the nucleolus, the subnuclear compartment where Pol I transcribes 45S pre-rRNA, but nothing is known about how rDNA copy number (dosage) affects ribosome biogenesis in muscle. Furthermore, while ribosome biogenesis is implicated in the muscle hypertrophic process, little is known about its contribution to the endurance exercise response.

We recently showed that an acute hypertrophic stimulus results in widespread CpG hypomethylation in promoter sites of genes related to growth (i.e. mTOR pathway and *Myc*) specifically within muscle fiber nuclei (myonuclei) (Von Walden *et al*., 2020a). These data concur with earlier findings showing epigenetic and signaling-related regulation of rDNA promoter regions in response to hypertrophic stimuli *in vivo* and *in vitro* in mice (von Walden et al., 2012; von Walden et al., 2016). Thus, early dynamic epigenetic events associate with robust transcriptional responses required for successful adaptation to exercise (Jozsi *et al*., 2000; Pilegaard *et al*., 2000). rDNA is highly regulated at the epigenetic level in general (Grummt, 2007; Murayama *et al*., 2008), but it is currently unknown whether rDNA promoter hypomethylation contributes to rRNA accumulation in response to exercise. Furthermore, epigenetic patterning and regulation of rDNA transcription is unique in part due to its tandem repeat organization and large intergenic spacer (IGS) (Baldridge *et al*., 1992; Mougey *et al*., 1996; Zentner *et al*., 2011a; Audas *et al*., 2012; Shiue *et al*., 2014), but there is no information on whether methylation of alternative regulatory sites in the rDNA repeat, such as enhancer regions and non-canonical transcription factor binding areas, are affected by mechanical loading and associate with ribosome biogenesis in muscle.

In the current investigation, we present the first time-course of ribosome biogenesis and rRNA transcription regulatory factor responses to acute endurance and resistance exercise (EE and RE, respectively) in human skeletal muscle, and comprehensively evaluate how rDNA gene dosage and methylation relates to rDNA transcription. To accomplish this, we employed whole-genome sequencing, targeted mass spectrometry-based rDNA promoter methylation analysis, reduced-representation bisulfite sequencing (RRBS), and mRNA- and protein-level measures. We complemented the human analyses with an analogous murine model of acute muscle loading, *in vivo* metabolic RNA labeling, and myonuclear-specific RRBS. With these parallel approaches, we reveal novel information on genetic and epigenetic transcriptional control of the translational apparatus following exercise. Our findings may have implications for individual heterogeneity in exercise responsiveness to training (Ahtiainen *et al*., 2016; Sparks, 2017; Lavin *et al*., 2019), as well as the epigenetic mechanisms of potentiated exercise adaptability related to long-term cellular “muscle memory” (Murach *et al*., 2019; Murach *et al*., 2020; Snijders *et al*., 2020).

## METHODS

### Research participants

Thirty healthy male and female participants (18 males and 12 females) were recruited and randomized to either a control (CON, n = 10), an endurance exercise (EE, n = 10) or a resistance exercise group (RE, n = 10). All groups included six males and four females. Participant characteristics are presented in Table 1. All participants were recreationally active, i.e. involved in EE one to three times per week and/or RE one to two times per week. Inclusion criterion was 18-50 years of age, and exclusion criteria were cardiovascular disease, neuromuscular disease, or severe knee problems. The study protocol was approved by the Regional Ethical Review board in Linköping and conformed to the Declaration of Helsinki. After receiving written and oral information about the study, the participants gave their informed consent to participate.

**Table 1 –.**
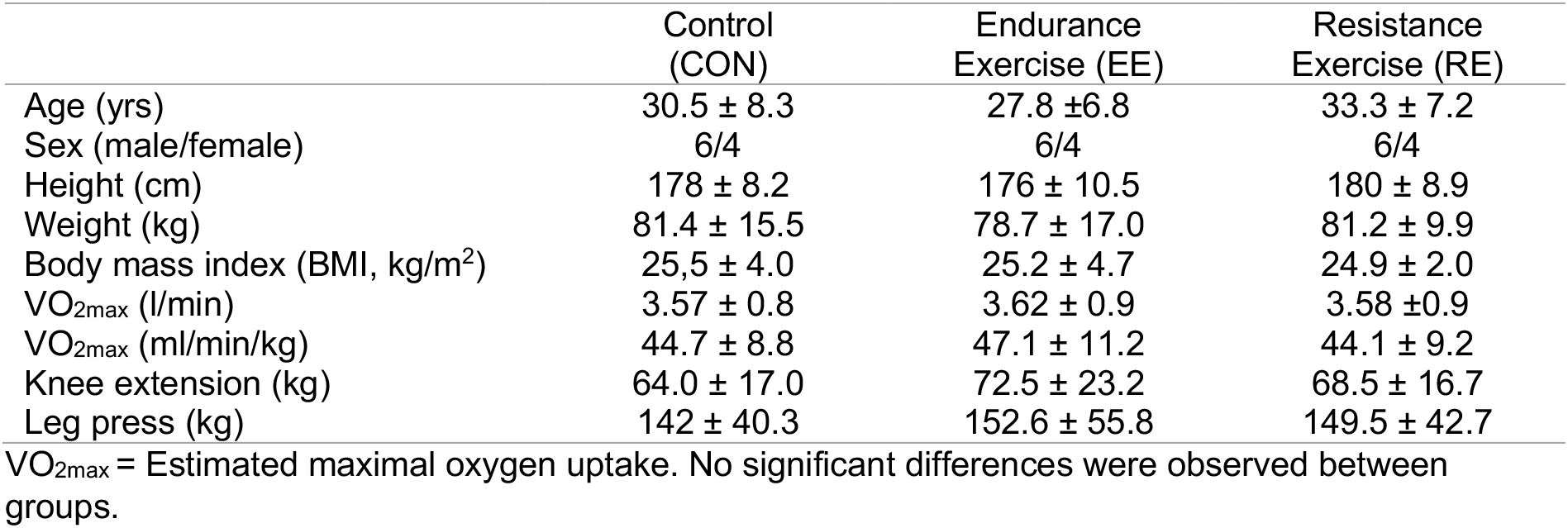
Characteristics of research subjects

### Human study design

At least five days prior to the intervention, participants were familiarized with the experimental set-up. All subjects performed a submaximal test on a cycle egometer (Monark 828 E, Monark Exercise AB, Vansbro, Sweden), to estimate VO_2_max (Ekblom-Bak *et al*., 2014; Bjorkman *et al*., 2016) and seven repetition maximum (7RM) for knee extension and leg press was titrated. These data were used to determine the load for the acute exercise bouts and to characterize the three groups with respect to physical status. Subjects were instructed not to perform any strenuous resistance for the legs three days prior to the intervention and no training the day prior. A liquid formula (1.05 g carbohydrates/kg body weight (bw), 0.28 g protein/kg bw, and 0.25 g fat/kg bw) was provided as breakfast 1h prior to collection of the pre-exercise sample and as lunch (2.10 g carbohydrates/kg body weight (bw), 0.56 g protein/kg bw, and 0.50 g fat/kg bw) immediately after the 3h biopsy and at breakfast on day 2, 2h before the final biopsy. The liquid formulas contained 5.6 g protein, 21 g carbohydrates, and 5.0 g fat per 100 ml (Resource komplett Näring, Nestlé Health Science, Stockholm, Sweden). Subjects were instructed to eat a standard dinner (plate model) the evening before the experiment and in the evening on the day of the experiment (Camelon *et al*., 1998). Skeletal muscle biopsies were collected before the intervention and at 30 min, 3h, 8h and 24h post exercise (Fig 1A). Eight subjects had 1-2 less biopsies taken due to various reasons (e.g. failed biopsy, adverse event) which affected the sample size for some analyses, and in some cases, there was only enough material for a specific analysis. Muscle biopsies were obtained from the vastus lateralis muscle percutaneously after injection of local anesthetic (carbocain 10 mg/ml), by using the Bergström biopsy needle with a diameter of 5 mm (Stille AB, Torshälla, Sweden). Skeletal muscle tissue was blotted for excess blood, cleaned of non-skeletal muscle tissue and snap-frozen in liquid nitrogen. All samples were stored at −80 °C until further analysis.

**Figure 1.**
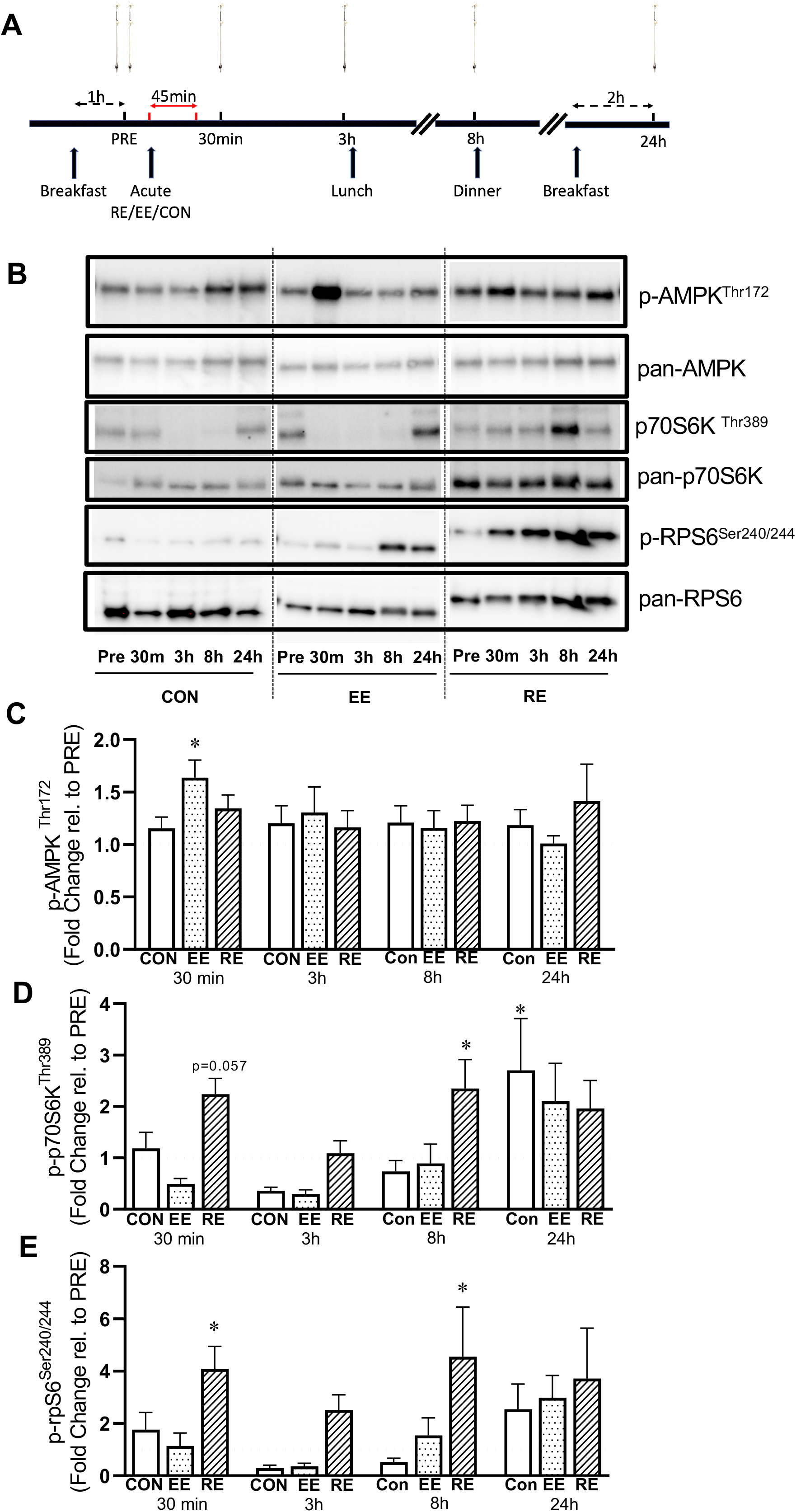
Exercise-related signaling in response to acute endurance exercise (EE) and resistance exercise (RE) over a 24-hour time course in human skeletal muscle **A.** Experimental study design timeline illustrating exercise and muscle biopsy collection in EE, RE, and control (CON) participants (n=10 per group) before (pre), 30 minutes, 3 hours, 8 hours, and 24 hours after exercise **B.** Western blot images for exercise-responsive protein targets in CON, EE, and RE **C.** Quantification of phosphorylated AMPK in CON, EE, and RE **D.** Quantification of phosphorylated p70S6K in CON, EE, and RE **E.** Quantification of phosphorylated rpS6 in CON, EE, and RE \**p*<0.05, Mean ± SE

### Human exercise protocols

RE consisted of two lower limb exercises; leg press (Nordic Gym AB, Bollnäs, Sweden) and knee extension (Nordic Gym AB). After a short warm-up on submaximal loads, the participants performed 4 sets per exercise at 7 repetition maximum load with 2 min rest between sets and 5 min between exercises. EE consisted of 45 min cycling (Monark 828 E, Monark Exercise AB, Vansbro, Sweden) at 70% of estimated VO_2max_. Heart rate was monitored continuously (Garmin Edge 25, Garmin, United States) and participants were asked to rate their level of perceived exertion every 5 min using the Borg RPE scale (Borg, 1970).

### Western blotting

Protein was extracted from the organic phase of TRI Reagent following RNA extraction using the optimized protocol (Wen *et al*., 2020). Briefly, following the final step of ethanol addition, samples were centrifuged and the resulting protein pellet solubilized using SDS-urea buffer (100 mM Tris, pH 6.8, 12% glycerol, 4% SDS, 0.008% bromophenol blue, 2% β-mercaptoethanol, 5 M urea) supplemented with Halt™ Protease (ThermoFisher #78438) and Phosphatase (#78426) Inhibitor Cocktails. RC DC™ Protein Assay (Bio-Rad, Hercules, CA) was used to determine protein concentration. Twenty micrograms of protein per sample was loaded on a gradient gel (Criterion™ Precast Gels, Bio-Rad) and electrophoretically transferred to a PVDF membrane (Bio-Rad). Pool control samples were loaded on all gels. Membranes were blocked in 5% bovine serum albumin (BSA, #A-420-1, Gold Biotechnology, St. Louis, MO), for phospho-specific antibodies, or 5% Non-Fat dry milk (#170-6404, Bio-Rad), for pan-antibodies, in Tris-buffered saline (TBS) with 0.1% Tween 20 (TBS-T) for two hours at room temperature. Following blocking, membranes were incubated overnight at 4°C with a primary antibody (all Cell Signaling Technology, Inc., Danvers, MA) in blocking solution. Antibodies used were: phospho-p70S6 Kinase (Thr389, 108D2, #9234, dilution 1:2000), pan-p70S6 Kinase (#9202, 1:2000), phospho-S6 Ribosomal Protein (Ser240/244, D68F8, #5364, dilution 1:2,000), pan-S6 Ribosomal Protein (5G10, #2217, dilution 1:3000), phospho-AMPKα (Thr172, 40H9, #2535, dilution 1:1000), pan-AMPKα (#2532, dilution 1:2000). The next day, membranes were incubated with a goat anti-rabbit (#G-21234, Thermo Fisher Scientific) secondary antibody (dilution 1:10,000 in blocking solution) for 1 hour at room temperature. Membranes were incubated with enhanced chemiluminescence (ECL) reagent (Clarity Western ECL substrate, #170-5060, Bio-Rad) before exposure to a ChemiDoc™ MP Imaging System (Bio-Rad). Bands were quantified using ImageJ software (NIH, Bethesda, MD). Coomassie blue staining was utilized to confirm equal loading.

### RNA extraction, cDNA synthesis and gene expression analysis

Approximately 25 mg of muscle tissue was used to extract RNA using TRI Reagent^®^ (Sigma-Aldrich, St. Louis, MO). Tissue was homogenized using beads and the Bullet Blender^®^ Tissue Homogenizer (Next Advance, Troy, NY). Following homogenization, RNA was isolated via phase separation by addition of bromochloropropane (BCP) and centrifugation. The supernatant was then transferred to a new tube and further processed on columns using the Direct-zol™ Kit (Zymo Research, Irvine, CA, USA). RNA was treated in-column with DNAse and eluted in nuclease-free water before being stored at −80°C. For quantitative reverse transcription PCR (qRT-PCR) 750 ng of total RNA was reverse transcribed using SuperScript™ IV VILO™ Master Mix (Invitrogen Carlsbad, CA). SsoAdvanced Universal SYBR Green Supermix (Bio-Rad, Hercules, CA) was used for quantitative reverse transcription PCR (qRT-PCR), on the CFX384 Thermocycler (Bio-Rad). PCR data was normalized by the geometric mean of three stable reference genes (*EMC7, VCP, C1ORF43*). Primer sequences are available upon request. Melting curves were performed for every primer pair to confirm a single-product amplification. qRT-PCR data were analyzed using the 2^−ΔΔ^CT method.

### Targeted human rDNA promoter methylation analysis

Genomic DNA was isolated from muscle samples using the QIAamp DNA Mini kit (Qiagen, Hilden, Germany) according to the manufacturer’s protocol. In brief, ~25mg frozen muscle samples were incubated and homogenized by enzyme digestion and mechanical disruption. After tissue had been completely dissolved, the mixture was added to mini spin-columns and a series of wash steps was performed. Each sample was diluted in 70 μl of distilled water. Immediately after DNA had been extracted, quantity and quality were determined in a NanoPhotometer NP80 (Implen, München, Germany).

Quantitative methylation analysis was performed using the EpiTYPER methodology (Ehrich et al. 2005) and the MassARRAY^®^ system (Agena Biosciences, San Diego, CA, USA) according to manufacturer’s recommendations and protocols, as previously described by our laboratory (von Walden *et al*. 2020b). In this method, a targeted amplification of bisulfite converted DNA is followed by *in vitro* transcription, RNase cleavage and subsequent fragment mass analysis by Matrix-Assisted Laser Desorption/Ionization Time of Flight Mass Spectrometry (MALDI-TOF MS) to quantify CpG sites. PCR primers were adapted from D’Aquilla et al. 2017 (D’Aquila *et al*., 2017). EpiTect methylated and non-methylated bisulfite-treated control DNA (Qiagen) was used to evaluate the quantitative recapture of methylation ratios of the amplicons. The amplicon used in this study met the quality criteria of methylated and non-methylated data points measured at > 79% and < 5% methylation ratios, respectively, as well as standard deviation percentages < 5%. Samples were run in duplicate and standard deviation percentages >20% were removed from the study (six out of 30 participants). The remaining data points (from n=24 participants) correlated with R^2^ 0.72. Bisulfite conversion efficiency was evaluated by analyzing one non-CpG C’s in a subset of the study samples. All data were checked by manually and visually inspecting the mass spectra.

### Estimation of rDNA copy number via quantitative PCR (qPCR)

Relative ribosomal DNA copy number was estimated by qPCR (rDNA dosage). Genomic (g)DNA was extracted from muscle biopsies (from n=27 participants) and isolated using the Monarch kit for DNA isolation (New England Biolabs, Ipswich, MA) according to the manufacturer’s instructions. Proteinase K digestion was performed overnight, and all samples were RNAse treated before being purified on column. gDNA was eluted in nuclease-free H_2_O and diluted to same concentration. 2.5 ng of DNA was loaded per well in triplicate. qPCR was run using Fast SYBR Green Master Mix (Applied Biosystems™, Foster City, CA) in a QuantStudio 3 Real-Time PCR Systems (Thermo Fisher Scientific, Waltham, MA). The sequence of the primers (18S, 5.8S and 28S, 5S and TP53 as reference gene), utilized in this study to assess rDNA dosage, are from (Gibbons *et al*., 2015): TP53 F 5’TGTCCTTCCTGGAGCGATCT3’ and R 5’CAAACCCCTGGTTTAGCACTTC3’; 5S rDNA F 5’TCGTCTGATCTCGGAAGCTAA3’ and R 5’AAGCCTACAGCACCCGGTAT3’; 5.8S rDNA F 5’CGACTCTTAGCGGTGGATCA3’ and R 5’GATCAATGTGTCCTGCAATTC3’; 18S rDNA F 5’GACTCAACACGGGAAACCTC3’ and R 5’AGACAAATCGCTCCACCAAC3’; 28S rDNA F 5’GCGGGTGGTAAACTCCATCT3’ and R 5’CACGCCCTCTTGAACTCTCT3’. Data were normalized to TP53 and expressed in arbitrary units (AU). Data were not compared to a standard curve with known rDNA quantity and is therefore referred to as relative rDNA dosage. Sample number is smaller (n=7) when comparing rDNA gene dosage to rDNA transcription at 24h post exercise due to technical reasons outlined above, and DNA for one sample not being suitable for analysis.

### Whole genome sequencing (WGS) and bioinformatics

Based on the qPCR results, DNA from the same samples from nine participants spread across the full range of rDNA gene dosage (n=3 low, n=3 middle, n=3 high) was selected for WGS. At least 1.5 μg of skeletal muscle DNA was used for analysis. To avoid potential amplification bias, a PCR-free protocol was used for library preparation. We calculated the rDNA copy number using a similar approach as previously described by Gibbons et al. 2014 by calculating relative depth differences between rRNA sequences and the background (whole genome). Reads were trimmed for adapter sequences and low quality (minimum phred score of 20) before aligning to the GRCh37 (hg19) human reference genome assembly using Bowtie2 v 2.3.4.3 with the “--end-to-end” option (Langmead & Salzberg, 2012; Langmead *et al*., 2019). Alignment results, produced in random order, were sorted with respect to their genomic positions using the samtools *sort* function and read depth at each position was computed using samtools *depth* function. We took advantage of the 45S rRNA sequence on the supercontig GL000220.1 in the GRCh37 (hg19) reference assembly and used this as a surrogate for the consensus rDNA repeat sequence (U13369), as described by Gibbons *et al*. We found that read depths computed using the supercontig rRNA regions were highly correlated with those computed from using U13369 (R^2^ > 0.96, data not shown). This approach precluded the need to generate a custom reference assembly to combine the rDNA sequence with the rest of the genome and ensured assembly version consistency, thereby limiting the confounding effects of the genome assembly version differences on the variability in background read depths among participants. Maximum read depth corresponding to each rRNA coding region (18S, 5.8S, and 28S) were divided by the average read depth for the whole genome to obtain rDNA component dosage, which is an estimate of the number of copies of rDNA in a haploid genome.

### Mouse study design

To specifically label myonuclei via genetic means, we generated female HSA^+/-^-GFP^+/-^ mice by crossing homozygous human skeletal actin reverse tetracycline transactivator (HSA-rtTA) mice developed by our laboratory (Iwata *et al*., 2018) with homozygous tetracycline response element histone 2B green fluorescent protein mice (TetO-H2B-GFP) obtained from the Jackson Laboratory (005104). GFP labeling is >90%, is highly specific to myonuclei, and does not result in labeling of satellite cell-derived myonuclei during the experimental period (Iwata *et al*., 2018), thus making the results specific to resident myonuclei. Mice were treated with doxycycline in drinking water (0.5 mg/ml with 2% sucrose) for one week. Following doxycycline treatment and a six day washout, mice underwent bilateral sham surgery (biological duplicate) or synergist ablation mechanical overload (OV) of the plantaris (biological triplicate) as described previously (von Walden *et al*., 2020a), then were euthanized 72 hours later. The mice in these experiments were ~3 months of age at the time of surgery, and immunohistochemistry and single fiber imaging for representative images is described in (von Walden *et al*., 2020a). For the metabolic RNA labeling experiments, age-matched C57BL/6J mice were subjected to sham and OV by the same surgeon as the HSA-rtTA mice (biological duplicate for sham and triplicate for OV).

### Mouse *in vivo* metabolic RNA labeling

Five hours prior to tissue harvesting after sham and OV, all mice were pulsed with 2 mg of 5-ethynyl-uridine (EU) dissolved in 200 μl of PBS via an intraperitoneal injection, as previously described by our laboratory (Kirby *et al*., 2016). Muscles were snap frozen in liquid nitrogen upon collection. RNA was extracted using TRIzol reagent (Invitrogen) and DirectZol columns with on column DNAse treatment (Zymo Research). RNA was resuspended in molecular-grade H_2_O and quantified by measuring the optical density (230, 260, and 280 nm) with a Nanodrop 1000 Spectrophotometer (ThermoFisher Scientific, Wilmington, DE). Nascent (EU-labeled) RNA was purified from 1 μg RNA per sample using the commercially available Click-iT Nascent RNA Capture Kit (Life Technologies, Carlsbad, CA). cDNA was generated on 500 ng of total RNA and all EU affinity-purified RNA using the SuperScript VILO cDNA Synthesis Kit (Life Technologies). We assessed relative gene expression in the total RNA and nascent RNA fractions by normalization to EMC7 by the comparative Ct (2^−ΔΔCt^) method.

### RRBS analysis of human skeletal muscle and mouse myonuclear DNA

DNA isolation from human biopsy samples (≤5 mg) and mouse myonuclear samples was carried out according to the detailed protocols of Begue et al. (Begue *et al*., 2017) and von Walden (Von Walden *et al*., 2020a), respectively. Briefly, using the QIAamp DNA micro kit (Qiagen), muscle samples or myonuclear pellets were re-suspended in “buffer ATL” supplemented with proteinase K overnight at 56°C. DNA binding to the column was conducted using 1 μg of carrier RNA (only for myonuclear samples), and washes and centrifugations were carried out according to the manufacturer’s instructions. DNA was eluted in 20 μl of molecular grade H_2_O and was stored at −80°C until the time of analysis. DNA quality assessment and RRBS was conducted in collaboration with Zymo Research. Quality and concentration were assessed using a Fragment Analyzer (AATI). “Classic” RRBS library preparation was performed by digesting 5 ng of genomic DNA with 30 units of MspI enzyme (New England BioLabs), and fragments were ligated to pre-annealed adapters containing 5’-methyl-cyotosine. Adapter-ligated fragments ≥50 bp were recovered using the DNA Clean & Concentrator Kit (Zymo Research, D4003) and bisulfite-treated using the EZ DNA Methylation-Lightning Kit (Zymo Research, D5030). Preparative-scale PCR was performed, and the products were purified again using the Clean and Concentrator kit. Paired-end sequencing was performed using Illumina HiSeq, and sequenced reads from bisulfite-treated libraries were identified using standard Illumina base calling software. Raw FASTQ files were adapter, filled-in nucleotides, and quality-trimmed using TrimGalore 0.6.4 using options for “--rrbs” and “--non_directional” mode retaining reads with minimum quality above phred score of 30. FastQC 0.11.8 was used to assess the effect of trimming and overall quality of the data. A custom genome assembly to interrogate rDNA methylation was generated by adding the consensus rDNA repeat sequences, GenBank U13369.1 (Gonzales and Sylvester, 1995) and BK000964.3 (Grozdanov *et al*., 2003), as a separate chromosome to the human (GRCh38p13) and mouse (GRCm39) reference genome assemblies, respectively. Due to mapping interference from highly similar sequences within the reference assemblies, these sequences were found using Blast by comparing the respective rDNA sequences to the reference assembly followed by masking of these sequences in the genome using N’s. Alignment to the custom human and mouse reference genomes was performed using Bismark 0.19.0. Methylated and unmethylated read totals for each CpG site were collected using the Methylation Extractor tool. Methylation levels of each sampled cytosine was estimated as the number of reads reporting a “C”, divided by the total number of reads reporting a “C” or “T”. Differential methylation analyses were performed using R Bioconductor package, methylSig v1.0.0 (Park *et al*., 2014), which accounts for both read coverage (minimum set to 5x, as previously described by Begue *et al*. 2017 and von Walden *et al*. 2020) and biological variation. Differentially methylated regions were determined with tiling using 100 bp segments and no minimum cutoff for CpG sites. The mouse data were analyzed using a beta-binomial distribution. The human data were analyzed using a generalized linear model accounting for the interaction between exercise status and time since the last bout and included 3 control participants in the model as a covariate (11 total samples), making for a robust statistical approach. CpG coverage in the human samples at individual sites can vary using RRBS, but in all instances were significant *p* values and low false discovery rate (FDR) was reported, CpGs in ≥5 participants were present in the dataset. The raw sequencing data were deposited in the NCBI Gene Expression Omnibus database (GSE144774).

Murine myonuclear isolations were conducted previously, as described by us (Von Walden *et al*., 2020a). Briefly, after euthanasia via lethal CO_2_ asphyxiation and cervical dislocation, plantaris muscles were harvested and processed for myonuclear isolation via fluorescent activated cell sorting (FACS). Muscle was dounce homogenized by hand in a sucrose-based physiological buffer, then the crude nuclear suspension was filtered through a 40 μm strainer and spiked with 4 μl of propidium iodide (PI, 1 mg/ml stock), then participanted to FACS analysis. Muscle from a control mouse (sucrose-only treated HSA-GFP) was used to determine background fluorescence, and myonuclei were classified as GFP+/PI+ after elimination of doublets via forward scatter area versus height (biological duplicate for sham and triplicate for overload) and collected in 15 ml conical tubes. Myonuclear suspensions were pelleted at 500 x g for five minutes using a swinging-bucket rotor prior to DNA isolation.

### Statistical analysis

Data was analyzed using a mixed-effects model in which exercise modality as one factor (EE vs. RE vs. CON) and time of biopsy as another (Pre, 30 min, 3h, 8h and 24h). Potential relationships among the variables assessed were investigated using one-tailed Pearson’s correlation analysis. When significant interactions or main effects were found, post hoc tests were performed using Sidak’s method. The level of significance was set at 5% (*p*< 0.05), and GraphPad Prism 9.0 software (San Diego, CA) was used for statistical analysis. For methylation data, *p* values <0.05 were utilized, and false discovery rate (FDR) is reported for differentially methylated CpG sites and regions in humans. Data are presented as means ± standard error of the mean, unless otherwise denoted.

## RESULTS

### Resistance exercise (RE) promotes mTORC1, while endurance exercise (EE) promotes AMPK signaling over the 24-hour time course of recovery in muscle

To verify that our chosen exercise modalities caused an increase in canonical signaling associated with exercise adaptation (see Figure 1A for study design), we analyzed the phosphorylation of key proteins involved with EE and RE (namely AMPK and mTORC1 signaling). As expected, EE and RE promoted different intramuscular signaling patterns. Only EE induced phosphorylation of AMPK at Thr172 30 minutes post exercise (*p*=0.012, Figure 1B & C). RE promoted signaling through mTORC1 as assessed by p70S6K at Thr389, specifically at 30 minutes (tendency, *p*=0.057) and 8 hours (*p*=0.042, Figure 1B, D, & E), while EE did not activate p70S6K (Figure 1D). In the control participants, p-p70S6K was induced at 24 hours after the pre biopsy following breakfast, indicating that food intake stimulates p-p70S6K to a similar extent as food intake combined with exercise 24 hours prior (*p*=0.038, Figures 2D). Similarly, phosphorylation of rpS6 at Ser240/244 increased mainly following RE, and no significant change was observed in CON or EE. At 30 min (*p*=0.0098) and 8h (*p*=0.0052), p-rpS6 was increased in the RE group compared to resting levels. Globally, EE stimulated AMPK while RE stimulated mTORC1 signaling, consistent with the literature (Vissing *et al*. 2013).

**Figure 2.**
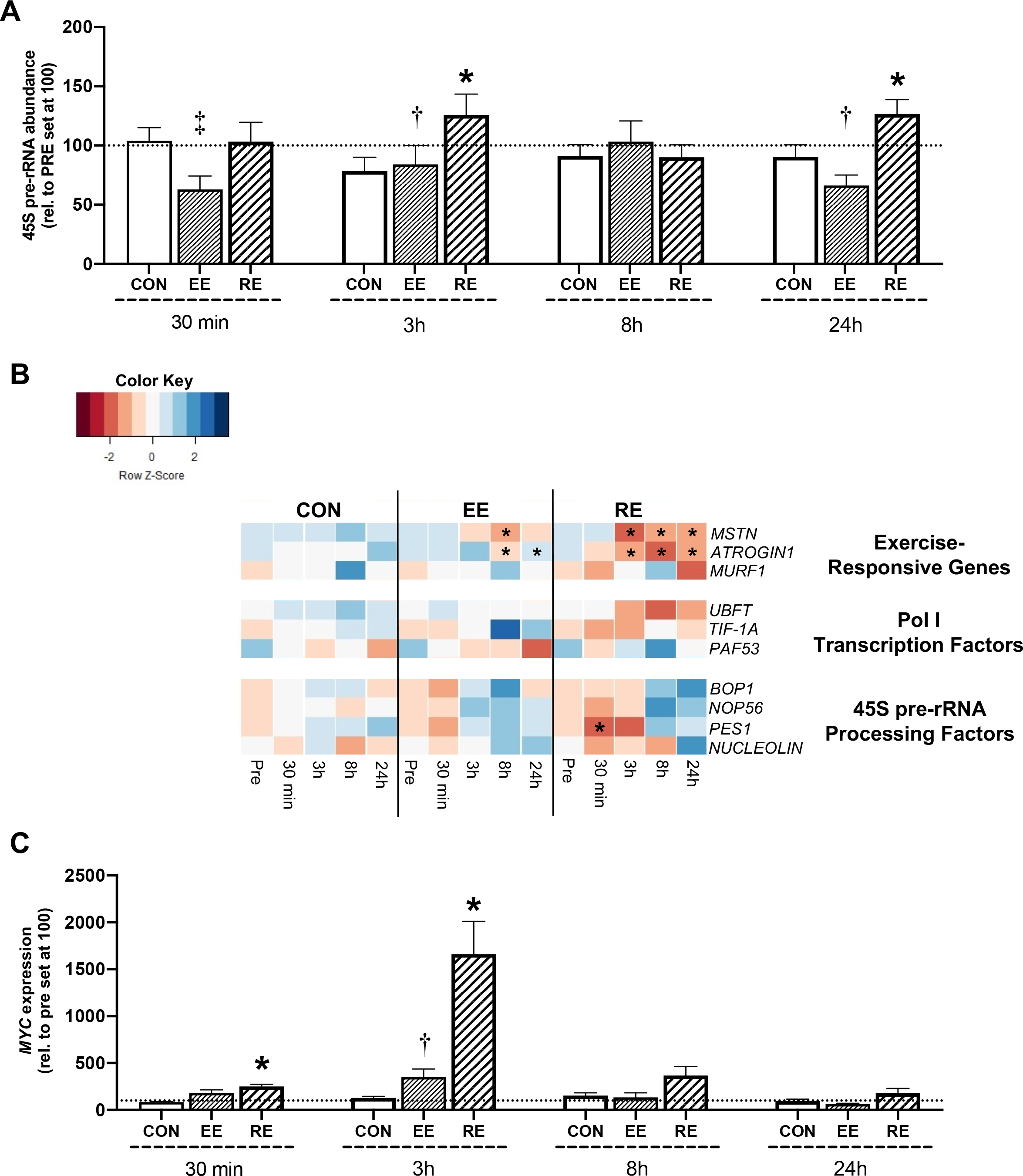
Transcriptional responses to acute endurance exercise (EE) and resistance exercise (RE) over a 24 hour time course in human skeletal muscle **A.** RNA Polymerase I (Pol I) transcription of ribosomal rDNA, as assessed by 45S pre-rRNA transcription levels, in control (CON), EE, and RE participants **B.** Heatmap illustrating gene expression for exercise-responsive genes (*MSTN, ATROGIN1, MURF1*), Pol I transcription factors (*UBFT, TIF-1A, PAF53*), and 45S pre-rRNA processing factors (*BOP1*, *NOP56, PES1, NUCLEOLIN*) in CON, EE, and RE participants **C.** *MYC* gene expression in CON, EE, and RE participants, normalized to Pre levels *&** *p*<0.05 & *p*<0.01 compared to CON, respectively; † *p*<0.05 compared to RE; ‡ *p*<0.05 compared to CON, Mean ± SE

### Resistance exercise preferentially stimulates ribosome biogenesis and Myc gene expression independent from transcription of RNA Pol I-associated factors

We utilized 45S pre-rRNA abundance as a readout of Pol I activity and ribosome biogenesis (Leary & Huang, 2001). EE caused an acute reduction of Pol I activity that was evident by 30 min post-exercise, and EE was significantly different than RE at both 3h and 24h (*p*<0.05, Figure 2A). Ribosome biogenesis was significantly induced by RE at 3h and 24h post-exercise, indicating an “early” and “late” ribosome biogenesis response. Genes that are responsive to exercise, *MSTN, ATROGIN1, and MURF1* (Louis *et al*. 2007), were generally affected in our time-course, but to a much greater extent with RE (*p*<0.01, Fig 2B). *MSTN* was reduced relative to CON 8h after EE and RE (*p*<0.05) and was also lower at 3h and 24h with RE (*p*<0.05). *ATROGIN1* was elevated at 3h and lower at 24h with EE (*p*<0.05), but was lower at 3h, 8h, and 24h with RE (*p*<0.05). There was an effect of time for *MURF1*, but otherwise no significant changes. Gene expression of factors associated with Pol I regulon formation and Pol I transcription (*UBTF, TIF1A, PAF53*) were not significantly affected by exercise, and therefore did not match the induction and suppression of 45S pre-rRNA seen following RE and EE, respectively (Figure 2B). Likewise, mRNA abundance of genes involved in rRNA processing (*BOP1, NOP56, PES1, and NUCLEOLIN*) were not significantly affected by acute exercise in either modality (Fig 2B), with the exception of a reduction in *PES1* 30 minutes after RE (p<0.05, Figure 2B). *C-MYC* (referred to as *MYC* for humans and *Myc* for mice), which is upstream of Pol I regulon formation (von Walden *et al*., 2012), was ~2 fold up-regulated in response to RE after 30 min and >15 fold higher after 3h (*p*<0.05, Figure 2C). Marked responsiveness of *MYC* within 24h of acute RE in vastus lateralis muscle is supported by meta-analytical data from resistance exercise transcriptome studies (Zierath and Pillon laboratories, www.MetaMEx.eu) spanning 110 healthy young and middle-aged men and women (log fold change=1.48, FDR=0.007×10^−4^) (Pillon *et al*. 2020); for reference, log fold change for *MSTN* in the same individuals was −0.81 (FDR=0.001×10^−4^). Expression levels of all genes in the control condition were not significantly altered at any time point for any gene.

### Relative rDNA gene dosage is related to rDNA transcription following a bout of resistance exercise

Since rDNA copy number can vary widely across people (Gibbons *et al*., 2014), we hypothesized that relative rDNA dosage would correlate with the ribosome biogenesis response to RE. First, to accurately assess relative rDNA gene dosage in our participants, we ranked all 30 individuals according to their normalized 18S rDNA gene content using a qPCR approach for rDNA quantification (Gibbons *et al*., 2014). The range of relative rDNA gene dosage among our 30 participants displayed a ~3-fold difference when comparing the lowest versus the highest values (Figure 3A). Based on this ranking, we selected the top three, bottom three, and three individuals in the middle of the distribution and performed whole genome sequencing (WGS) to directly assess rDNA copy number. From the WGS data, we calculated rDNA copy number and observed a significant positive correlation between this value and our qPCR quantification of 18S (r=0.793, *p*=0.005, Figure 3B) and 28S (*r*=0.590, *p*=0.047, Figure 3C) DNA. Moreover, the qPCR quantification of the different coding regions of the rDNA gene showed strong inter-correlation (28S versus 18S, r=0.846, *p*<0.0001, 28S versus 5.8S, r=0.975, *p*<0.0001, and 18S versus 5.8S, r=0.862, *p*<0.0001, Fig 3D-F), whereas quantification of the 5S rDNA gene (not part of 45S RNA gene) was less correlated with the other rRNA regions (5S versus 28S, r=0.297, *p*=0.066, 5S versus 18S, r=0.386, *p*=0.023, 5S versus 5.8S, r=0.382, *p*=0.042, Figure 3G-I). Relative rDNA dosage did not correlate to ribosome biogenesis at rest (data not shown). Finally, the increase in 45S pre-rRNA 24 hours after RE was positively correlated with relative rDNA dosage (18S, r=0.652, *p*=0.056; 28S, r=0.729, *p*=0.0313; 5.8S, r=0.696, *p*=0.041, Figures 3J-L).

**Figure 3.**
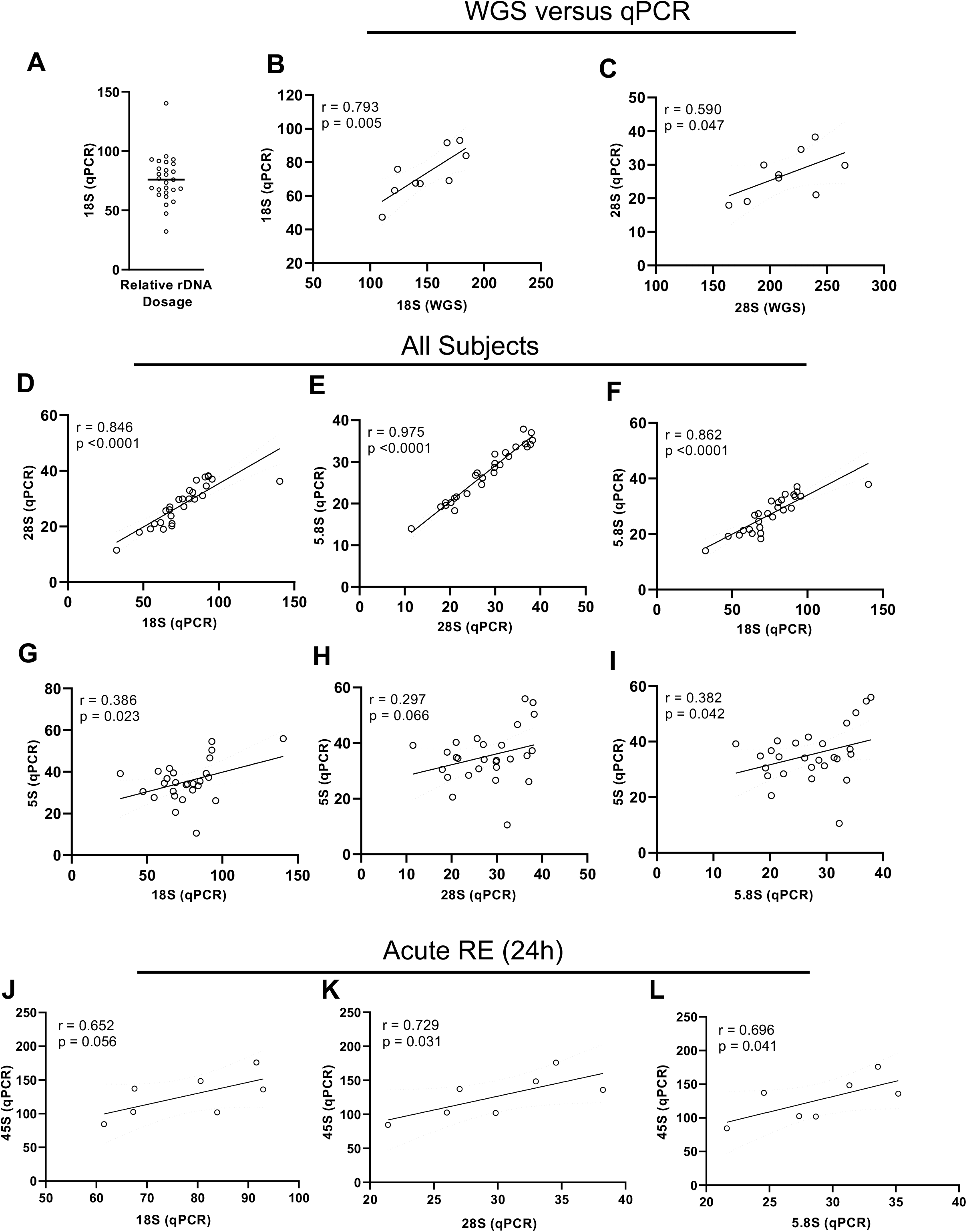
rDNA dosage for all participants at rest and its association to ribosome biogenesis in response to acute resistance exercise (RE) **A.** Relative rDNA dosage determined via q-PCR for all participants Comparison of rDNA copy number determined by whole genome sequencing (WGS) relative to rDNA copy number estimated by quantitative PCR (qPCR) using **B.** 18S rDNA and **C.** 28S rDNA Using the qPCR approach for relative DNA dosage determination, relationships between different locations in the 45S gene: **D.** 28S versus 18S, **E.** 5.8S versus 28S, and **F.** 5.8S versus 18S, as well as relationships between **G.** 5S (not part of 45S) and 18S **H.** 5S and 28S, and **I.** 5S and 5.8S Relationships between ribosome biogenesis (measured as 45S pre-rRNA) and estimated relative rDNA dosage using **J.** 18S, **K.** 28S, and **L.** 5.8S determined using qPCR

### Promoter methylation is not associated with ribosome biogenesis at rest

The human rDNA promoter encompasses a 174 base pair long section of the 45S RNA gene that include two important elements; the core promotor spanning −45 to +18 and the Upstream core element (UCE) −156 to −107, both relative to the transcription start site (Haltiner *et al*., 1986) (Figure 4A). We first used the Agena EpiTYPER, a sensitive and targeted mass spectrometry-based array method to compare the degree of methylation of the rDNA promoter in resting skeletal muscle at five sites within the promoter region. We calculated an average skeletal muscle rDNA promoter methylation of 23±8% in resting skeletal muscle (Figure 4B), in agreement with our previous work (Von Walden *et al*., 2020b), and consistent with levels reported in other human non-muscle tissues (Pietrzak *et al*., 2011; Uemura *et al*., 2012). Ribosomal DNA promoter methylation varied from 12% to 30% depending on the site, but site-specific methylation on a person-by-person basis tracked well (colored dots), so the average value is reflective of methylation along the promoter (Figure 4C). Average methylation was neither related to 45S pre-rRNA levels (r=-0.17, *p*=0.41, Fig 4D) nor total RNA content (r=-0.14, *p*=0.26, Fig 4E) at rest. These data collectively suggest that the regulation of ribosome biogenesis in skeletal muscle under resting conditions is not influenced by rDNA promoter methylation nor gene dosage.

**Figure 4.**
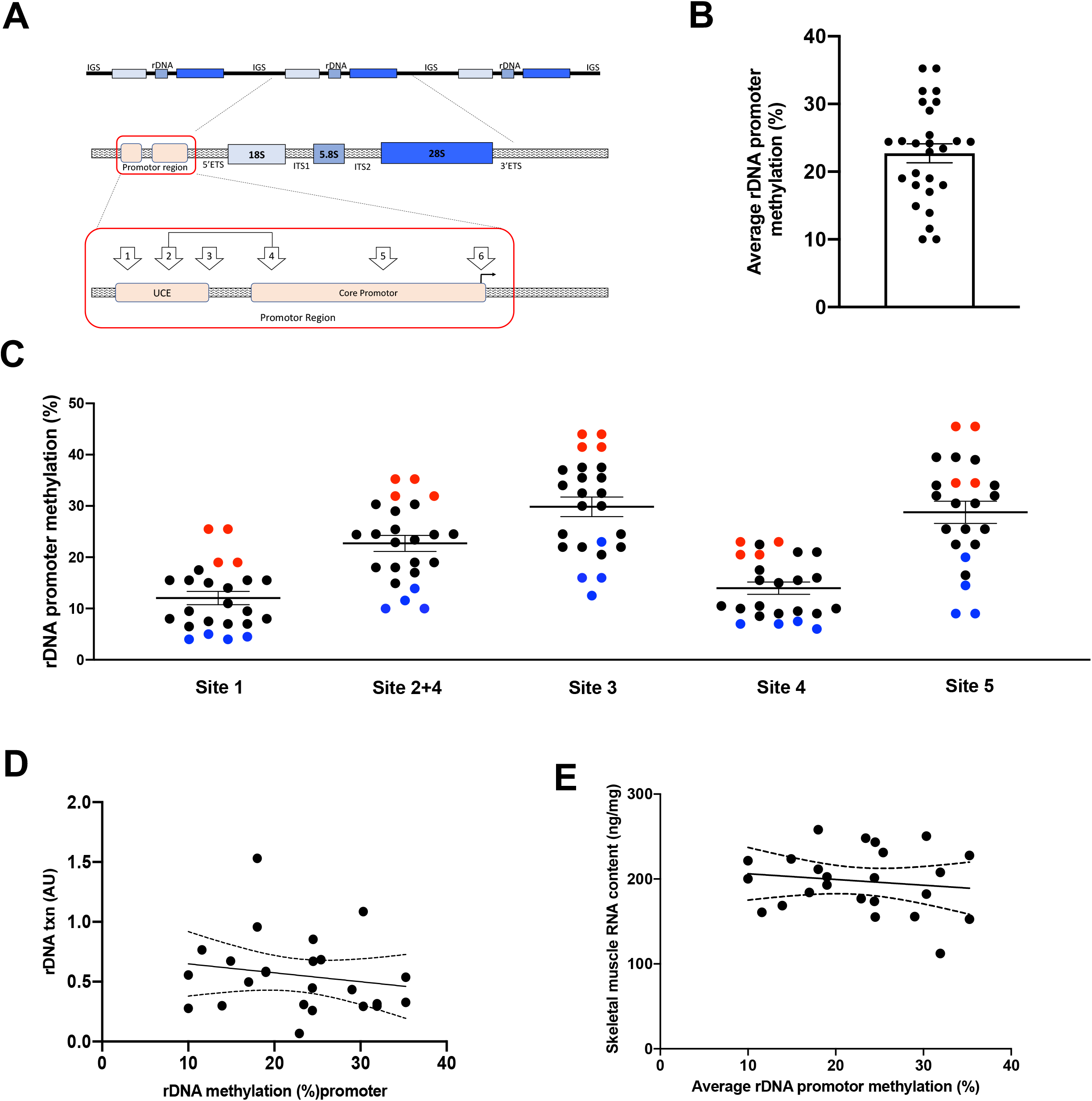
rDNA promoter methylation at rest measured via massARRAY EpiTYPER **A.** Illustration of specific CpG sites in the promoter region of rDNA that were measured in this study. Sites 2 and 4 were combined due to them having the same molecular weight **B.** Average rDNA promoter CpG methylation levels across all sites measured. Different colors represent the same participants across sites **C.** rDNA CpG methylation levels at individual sites in the promoter region **D.** Relationship between ribosome biogenesis and percent promoter methylation **E.** Relationship between RNA concentration (ng per mg of tissue) and percent CpG promoter methylation

### Resistance exercise acutely modifies methylation of the rDNA gene without affecting the promoter

To explore whether the acute modulation of 45S pre-rRNA abundance following EE (down-regulation) and RE (up-regulation) was associated with epigenetic modifications at the rDNA promoter, we analyzed targeted rDNA promoter methylation at all time points in all participants via EpiTYPER. We found no changes in average rDNA promoter methylation irrespective of exercise (Figure 5A). Although rDNA promoter methylation was not altered by exercise, the robust induction of ribosome biogenesis and *MYC* throughout the 24-hour time course following RE provided rationale to characterize methylation of other regulatory regions along the rDNA repeat in greater detail.

**Figure 5.**
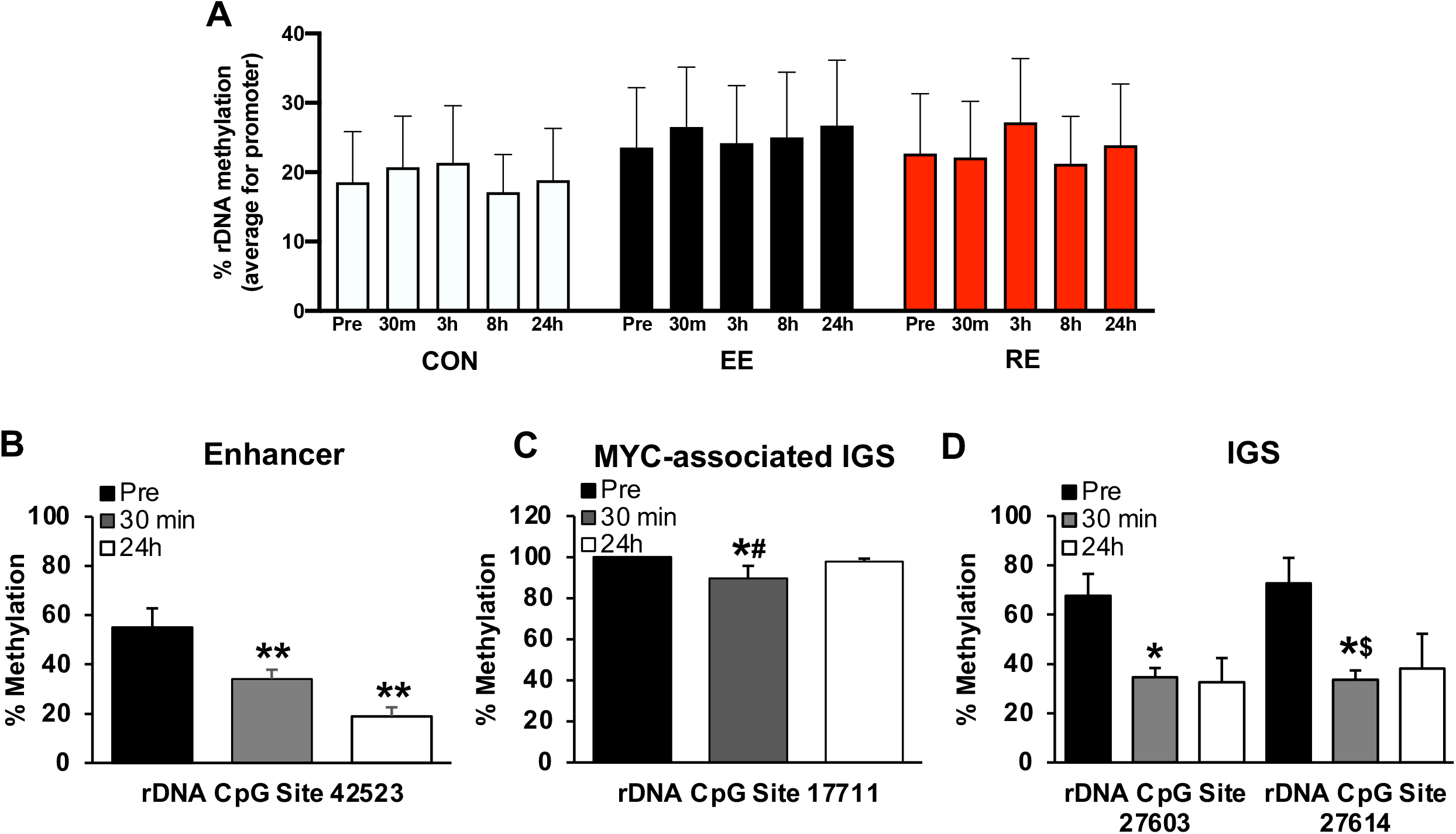
rDNA methylation in response to acute resistance exercise (RE) **A.** Promoter CpG methylation measured via massARRAY EpiTYPER in control (CON), EE, and RE participants over a 24-hour time course in human skeletal muscle **B.** CpG methylation at a site in the rDNA enhancer region pre, 30 minutes, and 24 hours after RE **C.** CpG methylation at a site in the rDNA associated with intergenic spacer (IGS) MYC occupancy pre, 30 minutes, and 24 hours after RE **D.** CpG methylation at sites in the IGS that is enhancer-like and immediately upstream of an IGS-derived transcript **FDR<0.05, ^#^FDR=0.05, \**p*<0.05, ^$^FDR=0.21

We conducted RRBS on DNA from a subset of RE participants at pre exercise, 30 minutes post, and 24 hours post exercise (n=8) and included data from 3 non-exercise participants at the corresponding time points in the analysis to account for the effects of biopsy and time. There is a strong interrelationship between chromatin modifications and DNA methylation (Eden *et al*., 1998; Fuks, 2005; Nemeth *et al*., 2008; Cedar & Bergman, 2009), so published chromatin immunoprecipitation (ChIP) and rDNA mapping data (Grandori *et al*., 2005; Zentner *et al*., 2011a; Shiue *et al*., 2014; Zentner *et al*., 2014; Agrawal & Ganley, 2018) were used to guide our analysis and infer implications of methylation patterns revealed by RRBS. Of the few CpG sites altered at both time points after RE, site 42523 relative to the transcription start site (TSS) had high coverage across individuals and was hypomethylated at 30 minutes (−17%, *p*=0.0002, FDR=0.03) and 24 hours (−32%, p=0.00003, FDR=0.01) (Figure 5B); this site is in a region characteristic of the rDNA enhancer (Zentner et al. 2011a), defined as H3K4me1/2 enriched (Zentner *et al*., 2011b). Since *Myc* mRNA was robustly up-regulated with RE, we looked for methylation differences in rDNA regions where MYC protein may associate. Thirty minutes after RE, an IGS CpG site in an area with MYC binding affinity (Grandori *et al*. 2005, Agrawal & Ganley, 2018) was hypomethylated (site 17711, −11%, *p*=0.0005, FDR=0.08) (Figure 5C). Immediately upstream of a region where MYC is most enriched on rDNA (~13 kb from the TSS) (Grandori *et al*. 2005), a CpG site was hypomethylated 30 minutes after RE (site 12054, 98% versus 88%, *p*=0.0007, FDR=0.09). Human rDNA has a unique enhancer-like region in the IGS (Zentner *et al*., 2011a) that contains MYC occupancy sites (Agrawal & Ganley, 2018) and codes for a stress-responsive IGS transcript (IGS_28_RNA) (Audas *et al*., 2012). Immediately upstream of this region, two highly methylated CpG sites in close proximity (site 27603 and 27614) were both ~30% hypomethylated 30 minutes after RE (*p*=0.0003, FDR=0.05 and *p*=0.002, FDR=0.21, respectively) (Figure 5D). Other hypomethylated CpG sites 30 minutes after RE were 4159 and 34987, and hypermethylated sites were 11938, 20819, 33349, and 33624 (FDR<0.05). 24 hours after RE, CpG sites 37512 and 39163 were hypomethylated, and 765, 11938, 20819, 33349, 33806, and 34428 were hypermethylated (FDR<0.05). Finally, a cluster of hypermethylated CpG sites 30 minutes after RE emerged 4-5 kb downstream of the TSS in the 18S coding region (4359, 4367, 4369, 4374, and 4382) (FDR<0.05).

### MYC transcription and rDNA methylation changes observed in human biopsy samples with RE were conserved with an acute hypertrophic stimulus in mice

Since skeletal muscle is a mixed tissue containing nuclei from various cell types, any epigenetic changes in rDNA genes in myonuclei could be obscured by non-muscle nuclei. Using a novel murine *in vivo* genetic myonuclear labelling strategy (HSA-GFP) to investigate global DNA methylation changes in response to an acute hypertrophic stimulus, we previously published that the responses are distinct between purified myonuclei and interstitial nuclei (von Walden *et al*., 2020a). Here, we confirm that 72 hours of mechanical overload (OV) of the plantaris muscle via synergist ablation in mice was associated with significantly higher abundance of the 45S pre-rRNA (Figure 6A). *In vivo* metabolic labeling of nascent RNA revealed that greater levels of rRNA was due to *de novo* synthesis, as illustrated by the elevated abundance of this transcript in the EU-labeled fraction following OV (Figure 6A). *Myc* was also highly enriched in the EU fraction after 72 hours of OV (Figure 6B). This finding verified that *Myc* is primarily regulated at the transcriptional level in response to OV, and not by post-transcriptional mechanisms, such as enhanced mRNA stability.

**Figure 6.**
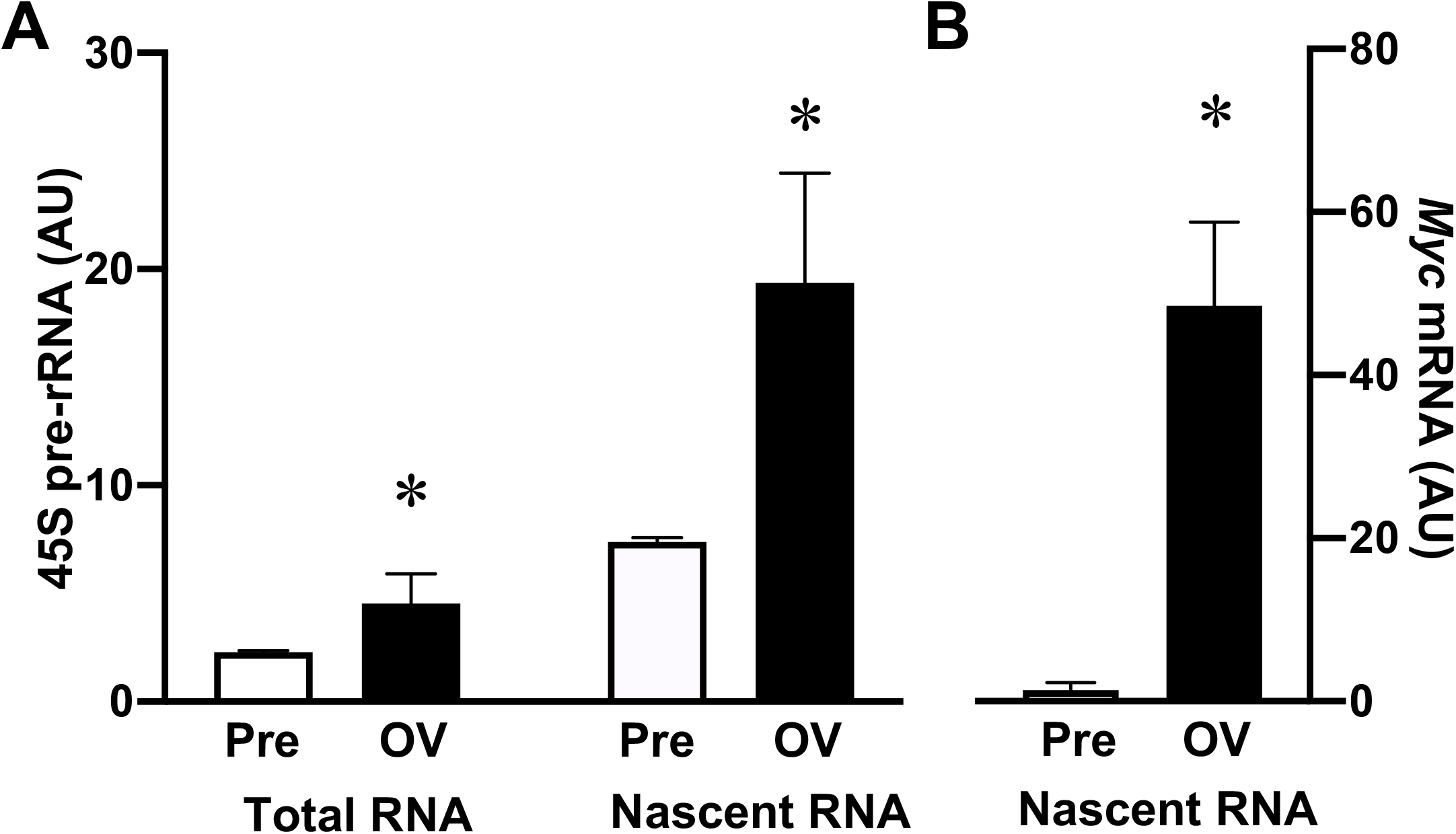
Ribosome biogenesis and *Myc* levels in acutely (72 hour) overloaded (OV) plantaris muscle with metabolic RNA labeling **A.** Total and metabolically labeled (nascent) rRNA in sham and 72 h OV muscle **B.** *Myc* levels in the nascent RNA fraction in sham versus OV muscle \**p*<0.05, Mean ± SE

To complement the human RRBS data, we gathered detailed information on the epigenetic regulation of rDNA specifically within myonuclei in response to an acute hypertrophic stimulus in mice. Labeled myonuclei from HSA-GFP mice were isolated via FACS (Figure 7A) and myonuclear rDNA methylation in sham and 72-hour OV plantaris muscle was analyzed (Figure 7B). DNA methylation patterns were altered in regions throughout the rDNA repeat. We observed differential methylation patterns with OV relative to sham in an rDNA enhancer region ~43-45 kb downstream of the TSS (Zentner *et al*., 2014), with hypermethylation in the 44501-44600 kb region (*p*=0.02, FDR=0.14) and hypomethylation in the 44701-48000 kb region (*p*=0.002, FDR=0.08) (Figure 7B). In this enhancer region, differential methylation generally occurred at sites with higher levels of methylation (>50%), and where methylation was >50% at either time point, hypomethylation predominated with OV (*p*≤0.05, Figure 7C). There are three major occupancy regions for Myc protein in mouse rDNA according to ChIP-sequencing: canonical binding in the upstream core element/promoter, a site within 1 kb downstream of the TSS, and in a ~7-13 kb downstream region (Zentner *et al*., 2014); differential methylation during OV coincides with the latter two regions (Figure 7A). The 201-300 kb (−19%) and 401-500 kb (−10%) regions downstream of the TSS were hypomethylated with OV (*p*=0.01, FDR=0.12 and *p*=0.02 and FDR=0.14, respectively), within which specific CpG sites were significantly hypomethylated (−19% at site 203 and −61% at site 466, *p*<0.05). The 11601-11700 downstream region was hypermethylated during OV (*p*=0.02, FDR=0.14) with an individual significantly hypermethylated CpG site therein (+26% at site 11674, *p*<0.05). On average, methylation in the 7-13 kb region was low and slightly hypermethylated with OV (~23% versus 27%), but we observed that in individual differentially methylated CpG sites where methylation was above 50% at either time point in this region, hypomethylation with OV predominated (*p*≤0.05, Figure 7D). Finally, in the IGS, the 39401-39500 region was hypomethylated after OV (−9%, *p*=0.03, FDR=0.14) with one specific CpG site therein trending to be hypomethylated with OV (−8% at 39484, *p*=0.06).

**Figure 7.**
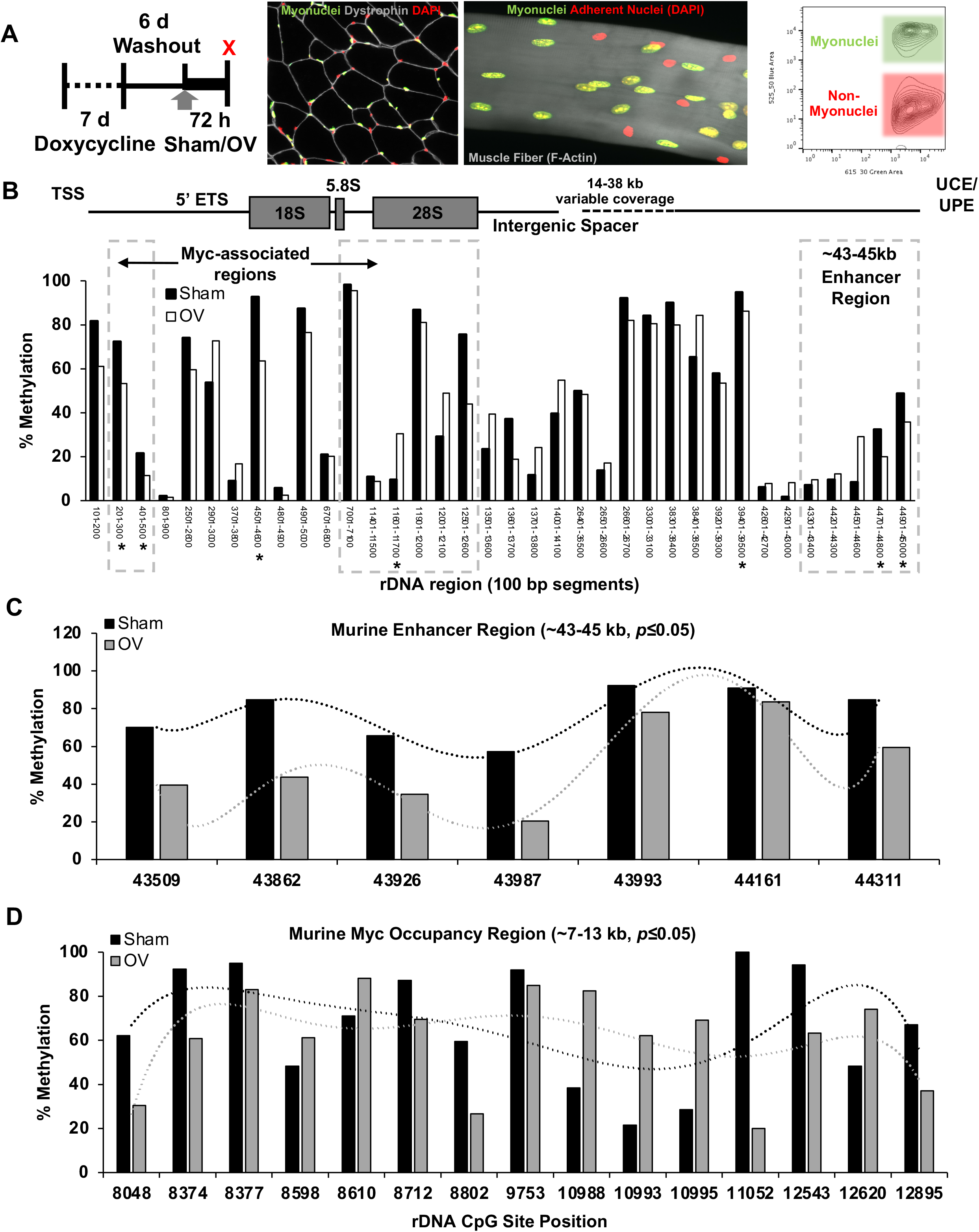
Myonuclear-specific rDNA methylation in acutely overloaded (OV) mice measured via reduced representation bisulfite sequencing (RRBS) **A.** Experimental study design illustrating the treatment and OV of HSA-GFP mice, myonuclear-specific GFP labeling, and isolation of highly-purified GFP+ myonuclei via fluorescent activated cell sorting **B.** Myonuclear rDNA CpG methylation in sham and 72-hour OV plantaris muscle. rDNA repeat is depicted above the graph based on CpG coverage. **C.** CpG methylation in the murine rDNA enhancer region (sites with >50% methylation) **D.** CpG methylation in murine rDNA *Myc* occupancy region (sites with >50% methylation) \**p*<0.05

## DISCUSSION

This study reveals that rDNA transcription in response EE or RE is not associated with changes in promoter methylation, but may be influenced by methylation within enhancer, IGS, and MYC binding regions specifically with RE. The human findings are corroborated by myonuclear-specific rDNA methylation after acute OV in mice, indicating a conserved response in murine muscle fibers. rDNA copy number correlates to the ribosome biogenesis response to RE, while the skeletal muscle response to EE seems less reliant on genetic and epigenetic factors regulating rDNA transcription. Ribosome biogenesis is suppressed in response to EE, indicating that rDNA transcription is a feature specific to hypertrophic adaptations, at least within the 24 hour time window after an acute bout.

To characterize our stimuli at the molecular level, we measured ribosome biogenesis and phosphorylation status of key proteins in signaling transduction pathways known to be responsive to exercise, specifically AMPK and mTORC1. Acute EE may repress ribosome biogenesis (Hansson et al. 2019), and ribosome biogenesis throughout our time course (pre-45S rRNA) was generally repressed by EE but was induced at 3 hours and 24 hours after RE. AMPK signaling was more prominent with EE versus RE, broadly consistent with the literature (Vissing *et al*. 2013, Murach & Bagley, 2016). mTORC1 is involved in muscle ribosome biogenesis (von Walden *et al*., 2016) and p70S6K Thr^389^ is a direct target of mTORC1, known to be highly phosphorylated in response to RE. Phosphorylation of this site was significantly increased in the RE group throughout the time course, but was also stimulated by feeding alone and endurance exercise with feeding at the 24 hour time point, uncoupled from ribosome biogenesis. Our ribosome biogenesis findings generally provide human *in vivo* support for a hypothesis where mTOR promotes Pol I transcription whereas AMPK inhibits it (von Walden, 2019), but also suggest mechanisms beyond signaling play a role in ribosome biogenesis with acute exercise. Furthermore, changes in transcriptional regulators involved in ribosome biogenesis and rRNA processing were unrelated to changes in 45S pre-rRNA at any time point, whereas the early induction of *MYC* did coincide with ribosome biogenesis. While robust up-regulation of *MYC* is consistent with its role in hypertrophic muscle adaptation (Chaillou *et al*., 2014; Wen *et al*., 2016; Figueiredo & McCarthy, 2019b; von Walden, 2019), the overall signaling and transcriptional findings motivated us to explore alternative regulatory mechanisms for ribosome biogenesis.

Skeletal muscle mass regulation has long been hypothesized to possess a strong genetic component (Seeman *et al*., 1996). While the search for genetic explanations of muscle hypertrophy have largely focused on normal protein-coding genes and gene polymorphisms (Ivey *et al*., 2000; Riechman *et al*., 2004; Charbonneau *et al*., 2008; Li *et al*., 2014), rDNA copy number has been overlooked. We first utilized WGS to quantify rDNA copy number in nine select individuals, then correlated the results to a qPCR method using validated primers from a previous study (Gibbons *et al*., 2015) to confirm the veracity of this approach in our samples. The qPCR method correlated well to the WGS data, supporting qPCR based relative rDNA gene dosage estimation. In our cohort, there was a three-fold difference between individuals with the lowest and the highest relative rDNA dosage, aligning with what has been previously reported (Gibbons *et al*., 2014b). The available evidence indicates that 45S pre-rRNA levels may peak at 24h after RE (Figueiredo & McCarthy, 2019a). Similar to previous observations, 45S pre-rRNA levels were significantly upregulated at the 24h time point in the RE group (Stec *et al*., 2015; Figueiredo *et al*., 2016), and changes in 45S pre-rRNA at 24h after RE were associated with rDNA copy number. An early ribosome biogenesis response to RE (within 3h) could therefore be related to signaling events, while the later response (24h) could be more influenced by the available template (i.e., number of rDNA copies), as opposed to the activation of anabolic signaling transduction pathways. Post-exercise anabolic signaling may therefore be permissive for early activation but not determinant for sustained upregulation of 45S pre-rRNA levels following an acute bout of RE, which points to a genetic predisposition for hypertrophic responsiveness based on rDNA gene dosage.

In muscle undergoing hypertrophy, histone modifications at the rDNA promoter coincide with increased rDNA transcription (von Walden *et al*., 2012). In light of this epigenetic regulation of rDNA during muscle cell growth, and in context with prior studies reporting genome-wide promoter methylation changes with acute exercise (Barres *et al*., 2012; Sharples *et al*., 2016; Seaborne *et al*., 2018; Sharples & Seaborne, 2019; Turner *et al*., 2019; Maasar *et al*. 2020), we hypothesized that CpG methylation changes to the rDNA promoter region may associate with the response to RE in muscle; however, this was not the case. rDNA promoter methylation in our adult muscle samples was low at rest (~20% on average), consistent with our previous report in children (von Walden *et al*., 2020b), suggesting the promoter is generally available for transcription in a CpG methylation context, and this does not change with acute EE or RE.

In addition to promoters, various regulatory elements such as enhancer regions and non-canonical transcription factor binding regions may also influence rDNA transcription (Zentner *et al*., 2011a; Shiue *et al*., 2014; Zentner *et al*., 2014). Hypermethylation of rDNA enhancer regions generally signifies an inactive gene (Brock & Bird, 1997; Stancheva *et al*., 1997). Hypomethylation of rDNA enhancer sites in humans and mice after a hypertrophic stimulus, as well as near a presumed IGS enhancer site/transcript coding region (Audas *et al*., 2012; Zentner et al., 2011a) suggests epigenetic remodeling of the rDNA repeat may support rDNA transcription in muscle fibers in response to exercise. MYC, a universal amplifier of expressed genes (Nie *et al*., 2012; Nie *et al*., 2020), is an rDNA-associated transcription factor that is central to ribosome biogenesis (Arabi *et al*., 2005), protein synthesis (Van Riggelen *et al*., 2010; West *et al*., 2016), and growth (Kim *et al*., 2000; Xiao *et al*., 2001; Zhong *et al*., 2006). MYC localizes in myonuclei during development and hypertrophy (Alway, 1997; Veal & Jackson, 1998), and its DNA binding extends beyond canonical E-box motifs (Allevato *et al*. 2017, Guo *et al*. 2014, Nie *et al*. 2012 and 2020) and is inhibited by CpG methylation (Prendergast *et al*., 1991; Perini *et al*., 2005). A site in the promoter of *Myc* is hypomethylated after 72 hours of OV in mouse myonuclei (von Walden *et al*., 2020a), which corresponds with *Myc* transcription observed here, and progressive Myc protein accumulation after RE in human and rodent muscle (Apró *et al*. 2013, Figueiredo *et al*. 2016, Ogasawara *et al*. 2016). We speculate that altered methylation of *MYC*-associated areas in rDNA is one component of a coordinated epigenetic regulatory network involving *MYC* upregulation, sustained ribosome biogenesis, and hypertrophic adaptation in skeletal muscle. There is also a functional link between dynamic MYC binding to IGS rDNA, rearrangement of nuclear architecture, and rDNA transcription specifically during growth in human cells (Shiue *et al*., 2014). The ramifications of rDNA methylation changes throughout the IGS during muscle hypertrophy in mice and humans deserve further investigation, especially in light of unique functional roles for transcripts originating from the rDNA IGS (Mayer *et al*., 2006; Santoro *et al*., 2010; Audas *et al*., 2012; Vacík *et al*., 2019). Further inquiry into the consequences and stability of repeat-wide rDNA methylation with exercise will broaden the emerging area of epigenetic regulation of rDNA transcription in skeletal muscle.

The induction of rDNA transcription induced by RE is positively associated with rDNA gene dosage in humans, suggesting that rDNA copy number may be a genetic determinant of muscle adaption in response to RE. Given the large variability of rDNA copy number across individuals, and that the muscle hypertrophy response to exercise is highly heterogeneous (Churchward-Venne *et al*., 2015; Ahtiainen *et al*., 2016), we propose that rDNA dosage is a potential genetic factor that relates to skeletal muscle hypertrophy induced by resistance training. Acute RE also promotes a reorganization of rDNA methylation patterns along the repeat. The specific consequences and lasting effects of this acute epigenetic remodeling of rDNA deserve further investigation, but if persistent over time, could contribute to a previously observed epigenetic “epi-memory” of prior exercise exposure that facilitates future muscle adaptability (Sharples *et al*., 2016; Seaborne *et al*., 2018; Moberg *et al*., 2020; Murach *et al*., 2020; Snijders *et al*., 2020).

## Author Contributions

VCF conducted experiments and data analysis, generated figures, and drafted the manuscript

YW conducted experiments and data analysis, generated figures, and assisted with manuscript preparation

BA coordinated the study, assisted with data collection, and provided resources

RF-G assisted with data collection and provided resources

JN assisted with data collection and provided resources

IV conducted experiments and data analysis

TV conducted experiments

CBM conducted experiments

GEZ assisted with manuscript preparation, figure preparation, and data analysis

CAP provided resources

JJM provided resources

KAM coordinated the study, conducted experiments and data analysis, generated figures and the graphical abstract, provided resources, and drafted the manuscript

FvW coordinated the study, conducted experiments and data analysis, generated figures, provided resources, and drafted the manuscript

All authors read, edited, and approved of the final manuscript

## Acknowledgements

The authors would like to thank the subjects for their extraordinary effort and participation. We further greatly acknowledge the help from staff at Höglandssjukhuset District Hospital in Eksjö where the human study took place, especially Annica Eriksson, Lena Norrbrand, Björn Otto, and the personnel at the Orthopaedic Department. We wish to thank Jennifer Strange of the University of Kentucky Flow Cytometry Core, as well as Dr. Keith Booher at Zymo Research. The graphical abstract was generated using BioRender.

## References

Agrawal S & Ganley AR. (2018). The conservation landscape of the human ribosomal RNA gene repeats. PloS One 13, e0207531.

Ahtiainen JP, Walker S, Peltonen H, Holviala J, Sillanpää E, Karavirta L, Sallinen J, Mikkola J, Valkeinen H & Mero A. (2016). Heterogeneity in resistance training-induced muscle strength and mass responses in men and women of different ages. Age 38, 10.

Alway SE. (1997). Overload-induced C-Myc oncoprotein is reduced in aged skeletal muscle. J Gerentol Ser A 52, B203–B211.

Allevato M, Bolotin E, Grossman M, Mane-Padros D, Sladek FM, & Martinez E. (2017). Sequence specific binding by MYC/MAX to low-affinity non-E-box motifs. PloS One 12, e0180147.

Apró W, Wang L, Pontén M, Blomstrand E & Sahlin K. Resistance exercise induced mTORC1 signaling is not impaired by subsequent endurance exercise in human skeletal muscle. (2013). Am J Physiol Endo Metab. 305, E22–32.

Arabi A, Wu S, Ridderstråle K, Bierhoff H, Shiue C, Fatyol K, Fahlén S, Hydbring P, Söderberg O & Grummt I. (2005). c-Myc associates with ribosomal DNA and activates RNA polymerase I transcription. Nat Cell Biol 7, 303–310.

Audas TE, Jacob MD & Lee S. (2012). Immobilization of proteins in the nucleolus by ribosomal intergenic spacer noncoding RNA. Mol Cell 45, 147–157.

Baldridge GD, Dalton MW & Fallon AM. (1992). Is higher-order structure conserved in eukaryotic ribosomal DNA intergenic spacers? J Mol Evo 35, 514–523.

Barres R, Yan J, Egan B, Treebak JT, Rasmussen M, Fritz T, Caidahl K, Krook A, O’Gorman DJ & Zierath JR. (2012). Acute exercise remodels promoter methylation in human skeletal muscle. Cell Metab 15, 405–411.

Begue G, Raue U, Jemiolo B & Trappe S. (2017). DNA methylation assessment from human slow-and fast-twitch skeletal muscle fibers. J Appl Physiol 122, 952–967.

Bjorkman F, Ekblom-Bak E, Ekblom O & Ekblom B. (2016). Validity of the revised Ekblom Bak cycle ergometer test in adults. Eur J Appl Physiol 116, 1627–1638.

Borg G. (1970). Perceived exertion as an indicator of somatic stress. Scand J Rehabil Med 2, 92–98.

Brock GJ & Bird A. (1997). Mosaic methylation of the repeat unit of the human ribosomal RNA genes. Human Molec Gene 6, 451–456.

Camelon KM, Hadell K, Jamsen PT, Ketonen KJ, Kohtamaki HM, Makimatilla S, Tormala ML & Valve RH. (1998). The Plate Model: a visual method of teaching meal planning. DAIS Project Group. Diabetes Atherosclerosis Intervention Study. J Am Diet Assoc 98, 1155–1158.

Cedar H & Bergman Y. (2009). Linking DNA methylation and histone modification: patterns and paradigms. Nat Rev Gene 10, 295–304.

Chaillou T, Kirby TJ & McCarthy JJ. (2014). Ribosome biogenesis: emerging evidence for a central role in the regulation of skeletal muscle mass. J Cell Physiol 229, 1584–1594.

Charbonneau DE, Hanson ED, Ludlow AT, Delmonico MJ, Hurley BF & Roth SM. (2008). ACE genotype and the muscle hypertrophic and strength responses to strength training. Med Sci Sports Exerc 40, 677–683.

Churchward-Venne TA, Tieland M, Verdijk LB, Leenders M, Dirks ML, de Groot LCPGM & van Loon LJC. (2015). There are no nonresponders to resistance-type exercise training inolder men and women. J Am Med Dir Assoc 16, 400–411.

D’Aquila P, Montesanto A, Mandala M, Garasto S, Mari V, Corsonello A, Bellizzi D & Passarino G. (2017). Methylation of the ribosomal RNA gene promoter is associated with aging and age-related decline. Aging Cell 16, 966–975.

Eden S, Hashimshony T, Keshet I, Cedar H & Thorne A. (1998). DNA methylation models histone acetylation. Nature 394, 842–842.

Ekblom-Bak E, Bjorkman F, Hellenius ML & Ekblom B. (2014). A new submaximal cycle ergometer test for prediction of VO2max. Scand J Med Sci Sports 24, 319–326.

Figueiredo VC. (2019a). Revisiting the roles of protein synthesis during skeletal muscle hypertrophy induced by exercise. Am J Physiol Reg Integr Comp Physiol 317, R709–R718.

Figueiredo VC, Caldow MK, Massie V, Markworth JF, Cameron-Smith D & Blazevich AJ. (2015). Ribosome biogenesis adaptation in resistance training-induced human skeletal muscle hypertrophy. Am J Physiol Endo Metab 30, E72–E83.

Figueiredo VC, Englund DA, Vechetti IJ, Jr., Alimov A, Peterson CA & McCarthy JJ. (2019). Phosphorylation of eukaryotic initiation factor 4E is dispensable for skeletal muscle hypertrophy. Am J Physio Cell Physiol 317, C1247–C1255.

Figueiredo VC & McCarthy JJ. (2019b). Regulation of ribosome biogenesis in skeletal muscle hypertrophy. Physiology 34, 30–42.

Figueiredo VC, Roberts LA, Markworth JF, Barnett MPG, Coombes JS, Raastad T, Peake JM & Cameron-Smith D. (2016). Impact of resistance exercise on ribosome biogenesis is acutely regulated by post-exercise recovery strategies. Physiol Rep 4.

Fuks F. (2005). DNA methylation and histone modifications: teaming up to silence genes. Curr Opin Gene Devel 15, 490–495.

Gibbons JG, Branco AT, Godinho SA, Yu S & Lemos B. (2015). Concerted copy number variation balances ribosomal DNA dosage in human and mouse genomes. Proc Nat Acad Sci 112, 2485–2490.

Gibbons JG, Branco AT, Yu S & Lemos B. (2014a). Ribosomal DNA copy number is coupled with gene expression variation and mitochondrial abundance in humans. Nat Comm 5, 1–12.

Gonzalez IL & Sylverster JE. Complete sequence of the 43-kb human ribosomal DNA repeat: analysis of the intergenic spacer. (1995). Genomics, 27, 320–328.

Grandori C, Gomez-Roman N, Felton-Edkins ZA, Ngouenet C, Galloway DA, Eisenman RN & White RJ. (2005). c-Myc binds to human ribosomal DNA and stimulates transcription of rRNA genes by RNA polymerase I. Nat Cell Biol 7, 311–318.

Grozdanov P, Georgiev O & Karagyozov L. (2003). Complete sequence of the 45-kb mouse ribosomal DNA repeat: analysis of the intergenic spacer. Genomics 82, 637–643.

Grummt I. (2007). Different epigenetic layers engage in complex crosstalk to define the epigenetic state of mammalian rRNA genes. Hum Mol Gene 16, R21–R27.

Guo J, Li T, Schipper J, Nilson KA, Fordjour FK, Cooper JJ, Gordan R, Price DH. (2014). Sequence specificity incompletely defines the genome-wide occupancy of Myc. Genome Biol 15, 482.

Haltiner MM, Smale ST & Tjian R. (1986). Two distinct promoter elements in the human rRNA gene identified by linker scanning mutagenesis. Mol Cell Biol 6, 227–235.

Hammarstrom D, Ofsteng S, Koll L, Hanestadhaugen M, Hollan I, Apro W, Whist JE, Blomstrand E, Ronnestad BR & Ellefsen S. (2020). Benefits of higher resistance-training volume are related to ribosome biogenesis. J Physiol 598, 543–565.

Hansson B, Olsen LA, Nicoll JX, von Walden F, Melin M, Strömberg A, Rullman E, Gustafsson T, Fry AC, Fernandez-Gonzalo R, Lundberg TR. (2019) Skeletal muscle signaling responses to resistance exercise of the elbow extensors are not compromised by a preceding bout of aerobic exercise. Am J Physiol Reg Comp Integr Physiol. 317, R83–92.

Ivey FM, Roth SM, Ferrell RE, Tracy BL, Lemmer JT, Hurlbut DE, Martel GF, Siegel EL, Fozard JL, Jeffrey Metter E, Fleg JL & Hurley BF. (2000). Effects of age, gender, and myostatin genotype on the hypertrophic response to heavy resistance strength training. J Gerentol Ser A 55, M641–648.

Iwata M, Englund DA, Wen Y, Dungan CM, Murach KA, Vechetti IJ, Mobley CB, Peterson CA & McCarthy JJ. (2018). A novel tetracycline-responsive transgenic mouse strain for skeletal muscle-specific gene expression. Skelet Muscle 8, 33.

Jozsi AC, Dupont-Versteegden EE, Taylor-Jones JM, Evans WJ, Trappe TA, Campbell WW & Peterson CA. (2000). Aged human muscle demonstrates an altered gene expression profile consistent with an impaired response to exercise. Mech Age Dev 120, 45–56.

Kim HG, Guo B & Nader GA. (2019). Regulation of ribosome biogenesis during skeletal muscle hypertrophy. Exerc Sport Sci Rev 47, 91–97.

Kim S, Li Q, Dang CV & Lee LA. (2000). Induction of ribosomal genes and hepatocyte hypertrophy by adenovirus-mediated expression of c-Myc in vivo. Proc Nat Acad Sci 97, 11198–11202.

Kirby TJ, Patel RM, McClintock TS, Dupont-Versteegden EE, Peterson CA & McCarthy JJ. (2016a). Myonuclear transcription is responsive to mechanical load and DNA content but uncoupled from cell size during hypertrophy. Mol Biol Cell 27, 788–798.

Langmead B & Salzberg SL. (2012). Fast gapped-read alignment with Bowtie 2. Nat Meth 9, 357.

Langmead B, Wilks C, Antonescu V & Charles R. (2019). Scaling read aligners to hundreds of threads on general-purpose processors. Bioinformatics 35, 421–432.

Lavin KM, Roberts BM, Fry CS, Moro T, Rasmussen BB & Bamman MM. (2019). The importance of resistance exercise training to combat neuromuscular aging. Physiology 34, 112–122.

Leary DJ & Huang S. (2001). Regulation of ribosome biogenesis within the nucleolus. FEBS Lett 509, 145–150.

Li X, Wang SJ, Tan SC, Chew PL, Liu L, Wang L, Wen L & Ma L. (2014). The A55T and K153R polymorphisms of MSTN gene are associated with the strength training-induced muscle hypertrophy among Han Chinese men. J Sports Sci 32, 883–891.

Louis E, Raue U, Yang Y, Jemiolo B, Trappe S. (2007). Time course of proteolytic, cytokine, and myostatin gene expression after acute exercise in human skeletal muscle. J Appl Physiol. 103, 1744–51.

Maasar MF, Turner DC, Gorski PP, Seaborne RA, Strauss JA, Shepherd SO, Cocks M, Pillon NJ, Zierath JR, Hulton AT, Drust B, Sharples AP. (2020). The methylome and comparative transcriptome after high intensity sprint exercise in human skeletal muscle. bioRxiv, https://doi.org/10.1101/2020.09.11.292805.

Malinovskaya EM, Ershova ES, Golimbet VE, Porokhovnik LN, Lyapunova NA, Kutsev SI, Veiko NN & Kostyuk SV. (2018). Copy number of human ribosomal genes with aging: unchanged mean, but narrowed range and decreased variance in elderly group. Front Gene 9, 306.

Mayer C, Schmitz K-M, Li J, Grummt I & Santoro R. (2006). Intergenic transcripts regulate the epigenetic state of rRNA genes. Mol Cell 22, 351–361.

McCarthy JJ & Murach KA. (2019). Anabolic and catabolic signaling pathways that regulate skeletal muscle mass. In Nutrition and Enhanced Sports Performance, pp. 275–290. Elsevier.

Moberg M, Lindholm ME, Reitzner SM, Ekblom B, Sundberg C-J & Psilander N. (2020). Exercise induces different molecular responses in trained and untrained human muscle. Med Sci Sport Exerc 52, 1679–1690.

Moss T. At the crossroads of growth control; making ribosomal RNA. (2004). Curr Op Gene Dev. 14, 210–7.

Mougey EB, Pape LK & Sollner-Webb B. (1996). Virtually the entire Xenopus laevis rDNA multikilobase intergenic spacer serves to stimulate polymerase I transcription. J Biol Chem 271, 27138–27145.

Murach KA & Bagley JR. (2016). Skeletal muscle hypertrophy with concurrent exercise training: Contrary evidence for an interference effect. Sport Med 46, 1029–1039.

Murach KA, Dungan CM, Dupont-Versteegden EE, McCarthy JJ & Peterson CA. (2019). “Muscle memory” not mediated by myonuclear number?: Secondary analysis of human detraining data. J Appl Physiol 127.

Murach KA, Mobley CB, Zdunek CJ, Frick KK, Jones SR, McCarthy JJ, Peterson CA & Dungan CM. (2020). Muscle memory: myonuclear accretion, maintenance, morphology, and miRNA levels with training and detraining in adult mice. J Cachex Sarc Muscle doi: https://doi.org/10.1002/jcsm.12617.

Murayama A, Ohmori K, Fujimura A, Minami H, Yasuzawa-Tanaka K, Kuroda T, Oie S, Daitoku H, Okuwaki M & Nagata K. (2008). Epigenetic control of rDNA loci in response to intracellular energy status. Cell 133, 627–639.

Nakada S, Ogasawara R, Kawada S, Maekawa T & Ishii N. (2016). Correlation between Ribosome Biogenesis and the Magnitude of Hypertrophy in Overloaded Skeletal Muscle. PloS One 11, e0147284–e0147284.

Nemeth A, Guibert S, Tiwari VK, Ohlsson R & Längst G. (2008). Epigenetic regulation of TTF-I-mediated promoter–terminator interactions of rRNA genes. EMBO J 27, 1255–1265.

Nie Z, Guo C, Das SK, Chow CC, Batchelor E, Jnr SSS & Levens D. (2020). Dissecting transcriptional amplification by MYC. eLife 9, e52483.

Nie Z, Hu G, Wei G, Cui K, Yamane A, Resch W, Wang R, Green DR, Tessarollo L & Casellas R. (2012). c-Myc is a universal amplifier of expressed genes in lymphocytes and embryonic stem cells. Cell 151, 68–79.

Ogasawara R, Fujita S, Hornberger TA, Kitaoka Y, Makanae Y, Nakazato K, Naokata I. The role of mTOR signalling in the regulation of skeletal muscle mass in a rodent model of resistance exercise. (2016). Sci Rep. 6, 31142.

Park Y, Figueroa ME, Rozek LS & Sartor MA. (2014). MethylSig: a whole genome DNA methylation analysis pipeline. Bioinformatics 30, 2414–2422.

Parks MM, Kurylo CM, Dass RA, Bojmar L, Lyden D, Vincent CT & Blanchard SC. (2018). Variant ribosomal RNA alleles are conserved and exhibit tissue-specific expression. Sci Adv 4, eaao0665.

Perini G, Diolaiti D, Porro A & Della Valle G. (2005). In vivo transcriptional regulation of N-Myc target genes is controlled by E-box methylation. Proc Nat Acad Sci 102, 12117–12122.

Pietrzak M, Rempala G, Nelson PT, Zheng J-J & Hetman M. (2011). Epigenetic silencing of nucleolar rRNA genes in Alzheimer’s disease. PloS One 6, e22585.

Pilegaard H, Ordway GA, Saltin B & Neufer PD. (2000). Transcriptional regulation of gene expression in human skeletal muscle during recovery from exercise. Am J Physiol Endo Metab 279, E806–E814.

Pillon NJ, Gabriel BM, Dollet L, Smith JAB, Puig LS, Botella J, Bishop DJ, Krook A, Zierath JR. (2020). Transcriptional profoiling of skeletal muscle adaptations to exericise and inactivity. Nat Comm 11, 1–15.

Prendergast GC, Lawe D & Ziff EB. (1991). Association of Myn, the murine homolog of max, with c-Myc stimulates methylation-sensitive DNA binding and ras cotransformation. Cell 65, 395–407.

Riechman SE, Balasekaran G, Roth SM & Ferrell RE. (2004). Association of interleukin-15 protein and interleukin-15 receptor genetic variation with resistance exercise training responses. J Appl Physiol 97, 2214–2219.

Santoro R, Schmitz KM, Sandoval J & Grummt I. (2010). Intergenic transcripts originating from a subclass of ribosomal DNA repeats silence ribosomal RNA genes in trans. EMBO Rep 11, 52–58.

Seaborne RA, Strauss J, Cocks M, Shepherd S, O’Brien TD, van Someren KA, Bell PG, Murgatroyd C, Morton JP, Stewart CE & Sharples AP. (2018). Human skeletal muscle possesses an epigenetic memory of hypertrophy. Sci Rep 8, 1898.

Seeman E, Hopper JL, Young NR, Formica C, Goss P & Tsalamandris C. (1996). Do genetic factors explain associations between muscle strength, lean mass, and bone density? A twin study. Am J Physiol 270, E320–327.

Sharples AP & Seaborne RA. (2019). Exercise and DNA methylation in skeletal muscle. In Sports, exercise, and nutritional genomics, pp. 211–229. Elsevier.

Sharples AP, Stewart CE & Seaborne RA. (2016). Does skeletal muscle have an ‘epi’-memory? The role of epigenetics in nutritional programming, metabolic disease, aging and exercise. Aging Cell 15, 603–616.

Shiue C-N, Nematollahi-Mahani A & Wright AP. (2014). Myc-induced anchorage of the rDNA IGS region to nucleolar matrix modulates growth-stimulated changes in higher-order rDNA architecture. Nuc Acid Res 42, 5505–5517.

Snijders T, Aussieker T, Holwerda A, Parise G, van Loon L & Verdijk LB. (2020). The concept of skeletal muscle memory: evidence from animal and human studies. Acta Physiol e13465.

Sparks LM. (2017). Exercise training response heterogeneity: physiological and molecular insights. Diabetologia 60, 2329–2336.

Stancheva I, Lucchini R, Koller T & Sogo JM. (1997). Chromatin structure and methylation of rat rRNA genes studied by formaldehyde fixation and psoralen cross-linking. Nuc Acid Res 25, 1727–1735.

Stec MJ, Kelly NA, Many GM, Windham ST, Tuggle SC & Bamman MM. (2016). Ribosome biogenesis may augment resistance training-induced myofiber hypertrophy and is required for myotube growth in vitro. Am J Physiol Endo Metab 8, E652–E661.

Stec MJ, Mayhew DL & Bamman MM. (2015). The effects of age and resistance loading on skeletal muscle ribosome biogenesis. J Appl Physiol 119, 851–857.

Turner DC, Seaborne RA & Sharples AP. (2019). Comparative transcriptome and methylome analysis in human skeletal muscle anabolism, hypertrophy and epigenetic memory. Sci Rep 9, 1–12.

Uemura M, Zheng Q, Koh CM, Nelson WG, Yegnasubramanian S & De Marzo AM. (2012). Overexpression of ribosomal RNA in prostate cancer is common but not linked to rDNA promoter hypomethylation. Oncogene 31, 1254–1263.

Vacík T, Kereïche S, Raška I, Cmarko D & Smirnov E. (2019). Life time of some RNA products of rDNA intergenic spacer in HeLa cells. Histochem Cell Biol 152, 271–280.

Van Riggelen J, Yetil A & Felsher DW. (2010). MYC as a regulator of ribosome biogenesis and protein synthesis. Nat Rev Cancer 10, 301.

Veal E & Jackson M. (1998). C-myc is expressed in mouse skeletal muscle nuclei during post-natal maturation. Int J Biochem Cell Biol 30, 811–821.

Vissing K, McGee SL, Farup J, Kjølhede T, Vendelbo MH, Jessen N. (2013). Differentiated mTOR but not AMPK signaling after strength vs endurance exercise in training-accustomed individuals. Scand J Med Sci Sport 23, 355–66.

von Walden F, Rea M, Mobley CB, Fondufe-Mittendorf Y, McCarthy JJ, Peterson CA & Murach KA. (2020a). The myonuclear DNA methylome in response to an acute hypertrophic stimulus. Epigenetics 15, 1151–1162.

von Walden F. (2019). Ribosome biogenesis in skeletal muscle: coordination of transcription and translation. J Appl Physiol 127, 591–598.

von Walden F, Casagrande V, Östlund Farrants A-K & Nader GA. (2012). Mechanical loading induces the expression of a Pol I regulon at the onset of skeletal muscle hypertrophy. Am J Physiol Cell Physiol 302, C1523–C1530.

von Walden F, Fernandez-Gonzalo R, Pingel J, McCarthy J, Stål P & Pontén E. (2020b). Epigenetic marks at the ribosomal DNA promoter in skeletal muscle are negatively associated with degree of impairment in cerebral palsy. Front Ped 8, 236.

von Walden F, Liu C, Aurigemma N & Nader GA. (2016). mTOR signaling regulates myotube hypertrophy by modulating protein synthesis, rDNA transcription and chromatin remodeling. Am J Physiol Cell Physiol, ajpcell.00144.02016-ajpcell.00144.02016.

Wen Y, Alimov AP & McCarthy JJ. (2016). Ribosome biogenesis is necessary for skeletal muscle hypertrophy. Exerc Sport Sci Rev 44, 110.

Wen Y, Vechetti Jr IJ, Valentino TR & McCarthy JJ. (2020). High-yield skeletal muscle protein recovery from TRIzol after RNA and DNA extraction. BioTechniques 69, 264–269.

West DW, Baehr LM, Marcotte GR, Chason CM, Tolento L, Gomes AV, Bodine SC & Baar K. (2016). Acute resistance exercise activates rapamycin-sensitive and-insensitive mechanisms that control translational activity and capacity in skeletal muscle. J Physiol 594, 453–468.

Xiao G, Mao S, Baumgarten G, Serrano J, Jordan MC, Roos KP, Fishbein MC & MacLellan WR. (2001). Inducible activation of c-Myc in adult myocardium in vivo provokes cardiac myocyte hypertrophy and reactivation of DNA synthesis. Circ Res 89, 1122–1129.

Zentner GE, Balow SA & Scacheri PC. (2014). Genomic characterization of the mouse ribosomal DNA locus. G3 4, 243–254.

Zentner GE, Saiakhova A, Manaenkov P, Adams MD & Scacheri PC. (2011a). Integrative genomic analysis of human ribosomal DNA. Nuc Acid Res 39, 4949–4960.

Zentner GE, Tesar PJ & Scacheri PC. (2011b). Epigenetic signatures distinguish multiple classes of enhancers with distinct cellular functions. Genome Res 21, 1273–1283.

Zhong W, Mao S, Tobis S, Angelis E, Jordan MC, Roos KP, Fishbein MC, de Alborán IM & MacLellan WR. (2006). Hypertrophic growth in cardiac myocytes is mediated by Myc through a Cyclin D2-dependent pathway. EMBO J 25, 3869–3879.

